# Ecological diversification in an adaptive radiation of plants: the role of de novo mutation and introgression

**DOI:** 10.1101/2023.11.01.565185

**Authors:** Benjamin W. Stone, Carolyn A. Wessinger

**Affiliations:** Department of Biological Sciences, University of South Carolina, Columbia, SC 29208-3401, USA

## Abstract

Adaptive radiations are characterized by rapid ecological diversification and speciation events, leading to fuzzy species boundaries between ecologically differentiated species. Adaptive radiations are therefore key systems for understanding how species are formed and maintained, including the role of de novo mutations vs. pre-existing variation in ecological adaptation and the genome-wide consequences of hybridization events. For example, adaptive introgression, where beneficial alleles are transferred between lineages through hybridization, may fuel diversification in adaptive radiations and facilitate adaptation to new environments. In this study, we employed whole-genome resequencing data to investigate the evolutionary origin of hummingbird-pollinated flowers and to characterize genome-wide patterns of phylogenetic discordance and introgression in *Penstemon* subgenus *Dasanthera*, a small and diverse adaptive radiation of plants. We found that magenta hummingbird-adapted flowers have apparently evolved twice from ancestral blue-violet bee-pollinated flowers within this radiation. These shifts in flower color are accompanied by a variety of inactivating mutations to a key anthocyanin pathway enzyme, suggesting that independent de novo loss-of-function mutations underlie parallel evolution of this trait. Although patterns of introgression and phylogenetic discordance were heterogenous across the genome, a strong effect of gene density suggests that, in general, natural selection opposes introgression and maintains genetic differentiation in gene-rich genomic regions. Our results highlight the importance of both de novo mutation and introgression as sources of evolutionary change and indicate a role for de novo mutation in driving parallel evolution in adaptive radiations.

## Introduction

Adaptive radiation – the proliferation of ecological roles and adaptations in different species within a lineage – is an important process considered to play a central role in the generation of biodiversity on Earth (Simpson 1953; Schluter 2000; Givnish 2015). This phenomenon, which often coincides with an increase in the rate of lineage diversification (and is thus termed a “rapid” or “explosive” adaptive radiation) encompasses a number of well-known and charismatic taxonomic groups, including Darwin’s finches, *Anolis* lizards, and cichlid fishes. Sustained interest in adaptive radiations from evolutionary biologists has, over time, revealed several genomic features characteristic of rapidly radiating clades. Principal among these is an abundance of phylogenetic discordance which, in depicting species’ complicated evolutionary histories, may more closely resemble a web than a bifurcating tree. This discordance may be chiefly caused by incomplete lineage sorting (ILS), which is expected to be abundant – potentially overwhelmingly so – when speciation occurs rapidly (Pamilo and Nei 1988; Maddison 1997; Rosenberg 2002; Whitfield and Lockhart 2007). Rapid adaptive radiations are also commonly characterized by introgression between closely related species that have not had much time to develop barriers to reproduction. Given that introgressed genetic material may confer fitness effects and can be subject to strong selection, particularly in functionally important genomic regions (Moran et al. 2021), signatures of introgression are expected to be distributed heterogeneously across the genome (Harrison and Larson 2016; Martin and Jiggins 2017). Introgression may provide a source of genetic diversity in adaptive radiations (Edelman and Mallet 2021), and although its importance in this context has historically been somewhat controversial (Seehausen 2004), many recent efforts have highlighted the myriad ways in which introgression may effectively broaden the genomic substrate for adaptation through novel combinations of haplotypes (e.g., McGee et al. 2020; Ferreira et al. 2021; Ronco et al. 2021; De-Kayne et al. 2022; Suvorov et al. 2022; Cicconardi et al. 2023; Marcionetti and Salamin 2023).

In addition to widespread ILS and introgression, radiations also often feature patterns of repeated evolution, which may be either the result of the adaptive introgression of beneficial alleles or independent (convergent) adaptation in disparate lineages to similar ecological conditions. While adaptive divergence progresses in adaptive radiations through complex interplays of standing genetic variation, introgression, and de novo mutation, it is standing or pre-existing variation (often introduced through introgression) that is often implicated as the most important source of genetic variation underlying repeated or parallel evolution (Berner and Salzburger 2015; Haenel et al. 2019; Marques et al. 2019; Marques et al. 2022). Yet, it is clear that the repeated generation of similar phenotypes in ecologically similar habitats – a hallmark of adaptive radiation – is also driven by de novo mutation. For example, both processes appear to be important in the radiation of hares (genus *Lepus*), in which the repeated evolution of nonwhite winter coats has proceeded through both adaptive introgression (Jones et al. 2018; Giska et al. 2019) and independent loss-of-function (LOF) mutations at the *Agouti* pigmentation gene (Jones et al. 2020). Whether adaptation more often involves de novo vs. standing variation informs on whether trait evolution is constrained by mutation events or by the opportunity for hybridization between differentiated lineages (Marques et al. 2019). Empirical data from diverse lineages across the tree of life are needed to identify common genetic features of adaptive radiations.

The genus *Penstemon* is an adaptive radiation of North American angiosperms which exhibits an astonishing array of phenotypic and ecological diversity across its nearly 300 described species. Many *Penstemon* species are interfertile and form hybrids in nature, fueling speculation that interspecific hybridization has played a major role in the diversification of the group (Straw 1955; Wolfe et al. 2006; Wessinger et al. 2016). *Penstemon* is also a recent and rapid radiation, estimated to have originated 1.419-6.416 MYA (Wolfe et al. 2021), and this rapid speciation has been highlighted as a potential major source of genealogical discordance (due to ILS) in addition to discordance caused by interspecific hybridization (Wolfe et al. 2006; Wessinger et al. 2016). Within *Penstemon*, the subgenus *Dasanthera* (hereafter, “*Dasanthera*”) includes nine ecologically diverse species and represents an adaptive radiation in microcosm. This group includes several species which hybridize in nature, making it an attractive system in which to study the importance of introgression and de novo mutation on adaptive divergence. A primary axis of variation in *Dasanthera* is floral pollination syndrome, wherein two hummingbird-adapted species (*P. newberryi* and *P. rupicola*) have evolved from the ancestral state of hymenopteran-adapted flowers (Datwyler and Wolfe 2004; Wilson et al. 2004; Kimball 2008). This mirrors a genus-wide trend involving the repeated evolution of specialist hummingbird-adapted flowers from more general hymenopteran-adapted progenitors (Wilson et al. 2007). The hummingbird-adapted species *P. newberryi* and *P. rupicola* each form hybrid zones with the hymenopteran-adapted, high-elevation *P. davidsonii*; in each case the hummingbird-adapted species occupies a lower elevation zone, and hybrids are formed and persist at intermediate elevations in the Cascades and Sierra Nevada Mountains where the parent species’ distributions overlap. Previous studies indicate that *P. newberryi* and *P. rupicola* are not sister species, suggesting parallel divergence in elevational tolerance and pollinator adaptation within *Dasanthera* (Stone and Wolfe 2021; Stone et al. 2023). An alternative scenario is that hummingbird pollination syndrome in *Dasanthera* could have a single evolutionary origin shared between species through introgression. Such a scenario has occurred in other plant groups; for example, the introgression of a *MaMyb2* allele has led to repeated origins of red-flowered phenotypes in the *Mimulus aurantiacus* species complex (Stankowski and Streisfeld 2015; Short and Streisfeld 2023).

Floral syndrome divergence in *Dasanthera* involves changes in floral morphology (hummingbird-pollinated: long, narrow flowers with exserted stamens and styles; hymenopteran-pollinated: short, wide flowers with inserted stamens and styles), and perhaps most strikingly, the evolution of bright magenta flowers in hummingbird-pollinated species from ancestral hymenopteran-pollinated blue-violet flowers. Whereas little is currently known about the underlying genetic mechanisms for floral morphological shifts, the evolution of red flowers tends to involve predictable changes to the genetic pathway for anthocyanin production. In other groups within *Penstemon*, shifts towards redder floral hues often involve the functional inactivation of the anthocyanin pathway enzyme Flavonoid 3′,5′-hydroxylase (F3′5′H) (Wessinger and Rausher 2014; Wessinger and Rausher 2015). F3′5′H belongs to the functionally diverse superfamily of cytochrome P450-dependent monooxygenases. This enzyme catalyzes hydroxylation of the B-ring of anthocyanins, leading to the production of blue-colored delphinidin-based pigments (Tanaka and Brugliera 2013). The degeneration of *F3′5′h* prohibits the production of delphinidin and has been shown in *Penstemon* to result in flowers that produce red pelargonidin-based anthocyanins; importantly, the evolution of red flowers in multiple *Penstemon* species appears to be the result of repeated de novo LOF mutations to the *F3′5′h* coding sequence (Wessinger and Rausher 2014; Wessinger and Rausher 2015).

In this study we used whole-genome resequencing data to trace the evolutionary history and source of genetic variation for hummingbird-pollinated flowers in *Dasanthera* and characterize genome-wide patterns of introgression. Signatures of introgression were uneven across the genome, and were generally lower in gene-rich regions, suggesting that even in this very recent radiation, introgression often may be maladaptive. We found strong support for independent evolutionary origins of magenta floral pigments in the two hummingbird-pollinated species. These parallel shifts in floral pigment production are accompanied by distinct degenerative mutations to the *F3′5′h* coding sequence. While introgression is prevalent within *Dasanthera*, we found little evidence that the two hummingbird-pollinated species share a history of introgression; this suggests that despite the importance of introgression across *Dasanthera* generally, it is de novo mutation, and not adaptive introgression, that likely fuels parallel shifts to hummingbird pollination in this adaptive radiation.

## Methods

### Study system and species sampling

We generated whole-genome resequencing data for 18 individual plants, each from a unique locality, representing seven of the nine total *Dasanthera* species (Table S.1). We included multiple samples for each of the species in the Cascades/Sierra Nevada Mountain clade (*P. cardwellii*, *P. davidsonii*, *P. fruticosus*, *P. newberryi*, *P. rupicola*), and a single individual each of the Northern Rocky Mountain exclusive taxa *P. montanus* and *P. lyallii* for use as outgroups, following results from previous phylogenetic inquiries into the group (Stone and Wolfe 2021; Stone et al. 2023). We extracted DNA from silica-dried leaf tissues using a modified CTAB protocol and performed whole-genome Illumina library preparations and sequencing to ∼10x depth using 150-bp paired-end reads generated on the NovaSeq (Illumina Inc.) platform at Duke University. Raw Illumina reads were quality-filtered with fastp (Chen et al. 2018), enabling auto-detection of adapters, limiting read length to 30 bp, filtering out unpaired reads, enabling base correction for overlapping reads, and enabling poly-x trimming on 3’ ends of reads. Final sequence quality was verified with fastqc (Andrews 2010) and multiqc (Ewels et al. 2016). Quality filtered reads were mapped to the annotated *P. davidsonii* reference genome (Ostevik et al. 2023) with bwa-mem (Li 2013), and reads with low mapping quality (q<20) were removed. Duplicated reads were marked and removed with samtools markdup (Li et al. 2009), and overlapping paired end reads were clipped with bamutil clipOverlap (Jun et al. 2015).

Filtered reads were then used to call genotypes to produce an “all sites” .vcf file using bcftools v1.15.1 (Li 2011). Variant sites were soft-filtered for minimum genotype-call quality (20), minimum and maximum allelic depths, which were chosen based on genome-wide averages of coverage depth (3 and 60, respectively), minimum individual allelic depth (2), and missingness (50%). Sites failing the minimum genotype-call quality filter were changed to match the reference allele (changed to invariant sites); otherwise, filters produced missing data for sites or individual genotypes. The resulting filtered all sites .vcf was then used to generate consensus whole-genome sequences and individual gene coding sequences (CDS).

### Species relationships

We inferred species trees in three ways to compare genomic sampling strategies and analysis methods. First, we used custom python scripts to split whole-genome sequences into non-overlapping 10kb windows, removing windows with more than 75% missing data (Ns) for any sample. We then estimated “window trees” for each genomic window with IQ-TREE 2 v2.2.0.3 (Nguyen et al. 2015), enabling the ModelFinder (-m MFP) option (Kalyaanamoorthy et al. 2017) to select the best substitution model for each window. We then used these window trees to infer a species tree with ASTRAL-III v.5.7.8 (Zhang et al. 2018), using the full annotation option (-t 2). Second, for each sample, we used gffread v0.12.7 (Pertea and Pertea 2020) and the *P. davidsonii* reference genome annotations to generate fasta files with spliced exons (CDS), retaining the single longest isoform for each transcript. We then used a custom python script to filter individuals with >50% missing data for each CDS. Gene trees were then estimated for each CDS and a species tree was constructed in ASTRAL in the same manner as for the window alignments. For both of the ASTRAL species trees, we also calculated gene-and site concordance factors (gCF, sCF; Minh et al. 2020) in IQ-TREE, using as input the window or gene trees and sequence alignments used to generate the species tree. Third, we estimated a concatenated ML species tree in IQ-TREE, specifying the GTR+I+R model, estimating rates for each of the 12 largest scaffolds, and performing 1000 UltraFast Bootstrap (UFBoot) replicates (Hoang et al. 2018).

### Floral pigment production

We determined the identity of anthocyanidin pigments produced in floral tissues, which are responsible for floral color in *Penstemon*. We collected intact flowers from voucher specimens that were preserved in an herbarium press at the time of collection (samples were collected in 2017 and 2018). We used an isoamyl alcohol extraction protocol outlined in Harborne (1984) and separated pigments using TLC on Cellulose F glass plates (Millipore, Burlington, MA). We compared extracted anthocyanidin pigments to a multi-anthocyanidin standard solution containing pelargonidin, cyanidin, and delphinidin each in concentrations of ∼0.013 mg/mL, as described in Stevens et al. (2023).

### Genome-wide average signatures of introgression

We investigated genome-wide signatures of introgression using Dsuite v0.5 (Malinsky et al. 2021). Specifically, we used Dsuite to calculate genome-wide D statistics (ABBA-BABA) for all possible trios of samples and to implement and visualize the “f-branch” metric *f_b_* (Malinsky et al. 2018) using the 10kb window ASTRAL species tree as the guide tree and *P. montanus* as the outgroup.

### Genomic landscape of divergence, discordance, and introgression

We examined patterns of differentiation among species and patterns of phylogenetic discordance across the genome. We calculated average *F_ST_* (Weir and Cockerham 1984) and average nucleotide distance within (*π*) and between (*d_xy_*) species in 10kb and 50kb non-overlapping windows with pixy v1.2.7 (Korunes and Samuk 2021), using the filtered all sites .vcf files as input. We used bedtools v2.27.1 (Quinlan and Hall 2010) to calculate genic fractions (the proportion of sites that are coding sequence) per window. We identified *d_xy_* outliers between two species pairs that form hybrid zones: *P. davidsonii*-*P. newberryi*, and *P. davidsonii*-*P. rupicola*. Highly differentiated genomic regions in the face of gene flow may be especially resistant to introgression and could play a key role in the formation or maintenance of species boundaries. We first filtered all sliding windows with more missing counts than count comparisons, and then identified outliers as 10kb sliding windows with average *d_xy_* values exceeding the genome-wide mean by three standard deviations (Z-score > 3). We then identified shared outliers as sliding windows with outlier *d_xy_* values in both species pairs.

To explore how topological discordance varies across the genome, we first calculated normalized Robinson-Foulds (RF) distances (Robinson and Foulds 1981) between each 10kb window tree and the species tree using the R package phangorn v2.10.0 (Schliep 2011). In addition, we examined topological discordance affecting three main internal branches of the species tree using TWISST (Martin and Van Belleghem 2017). TWISST quantifies the proportion (or “weighting”) of different tree topologies across the genome. For this approach, we used scripts provided by the developers (available at https://github.com/simonhmartin/twisst), using the 10kb window trees as input. We summed topology weights for all trees consistent with the following internal branches of the species tree: the branch leading to *P. newberryi*, *P. cardwellii*, and *P. rupicola* (termed IB1); the branch leading to *P. newberryi* and *P. cardwellii* (IB2), and the branch leading to *P. davidsonii* and *P. fruticosus* (IB3). In each case, *P. montanus* served as the outgroup.

Although the *D* statistic is a powerful metric to detect genome-wide patterns of introgression, it is less useful to identify specific introgressed genomic regions because it is a poor estimator of the amount of introgression and it has a high degree of variance when applied to genomic windows of smaller sizes (Martin et al. 2015). For this reason, to explore signatures of introgression along the genome, we employed the *f_dM_* statistic, which is specifically designed to estimate introgression between pairs of taxa in genomic windows along chromosomes, and, like many statistics related to the *f4*-ratio, performs better than the *D* statistic (exhibits less variance and is a better estimator of the amount of introgression) when applied to small genomic windows (Malinsky et al. 2015; Malinsky et al. 2021). We estimated *f_dM_* in 10,000 SNP windows, sliding every 2,500 SNPs. For each of these *f_dM_* windows, we also calculated the genic fraction to assess the relationship between genic fraction and *f_dM_*. We conducted these calculations for all triplets of taxa consistent with the species tree.

### Introgression of F3′5′h

We examined several lines of evidence to assess whether *F3′5′h* – one of the key genetic components underlying transitions to hummingbird pollination syndrome – could be shared between red-flowered *Dasanthera* through introgression. First, we calculated *f_dM_* in sliding windows along the genome to investigate signatures of introgression between *P. newberryi* (P2) and *P. rupicola* (P3), setting *P. cardwellii* as P1. Here we estimated *f_dM_* at four different window sizes: (1) in 10,000 SNP windows, sliding every 2,500 SNPs, (2) in 5,000 SNP windows, sliding every 1,250 SNPs, (3) in 1,000 SNP windows, sliding every 250 SNPs, and (4) in 500 SNP windows, sliding every 125 SNPs. If *F3′5′h* is introgressed, values of *f_dM_* should be elevated compared to both genome-wide estimates and the local genomic region.

Because *F3′5′h* is potentially pseudogenized in our magenta-flowered samples (leading to relaxed selection against indels), we re-called variants for reads mapping to this genomic region while additionally calling and normalizing indels with bcftools. We performed downstream variant calling and filtering as described for genome-wide variant calling steps. We then aligned the CDS for each sample with muscle v.5.1 (Edgar 2004) and re-estimated the gene tree for *F3′5′h* in IQ-TREE, performing 1000 UFBoot replicates and 1000 bootstrap replicates for the SH-like approximate likelihood ratio test (SH-aLRT) to assess branch support (Guindon et al. 2010). If *F3′5′h* is introgressed, this gene tree should be discordant with the species tree and place *P. newberryi* and *P. rupicola* as sister.

We examined *F3′5′h* nucleotide and amino acid sequence alignments to identify the positions of potential indels and nonsynonymous mutations. *F3’5’h* coding sequences in other hummingbird-adapted *Penstemon* species exhibit a variety of functional inactivating mutations, including premature stop codon mutations, frameshift mutations, and large deletions (Wessinger and Rausher 2015). We searched for similar LOF mutations present in our magenta-flowered samples. Likewise, mutations in any of the highly conserved domains, which included the heme-binding domain and six substrate recognition sites (SRSs) putatively involved in substrate contact (Gotoh 1992), would likely signal a dysfunctional enzyme. If *F3′5′h* is introgressed, our magenta-flowered samples should share the same loss-of-function mutation(s), be they through structural changes (e.g., indels) or through mutations in conserved domains or SRSs.

## Results

### Whole genome data confirm species relationships and suggest two separate origins of hummingbird pollination in Dasanthera

Our whole genome sequencing of 18 *Dasanthera* samples produced ∼13x average coverage per sample across approximately 70% of the eight main scaffolds (assumed to represent the eight *Penstemon* chromosomes). All information pertaining to read quantities, genome-wide coverage, and per-scaffold coverage can be found in Table S.2.

All three species tree inference methods produced very similar topologies and support the same main relationships among species within the Cascades and Sierra Nevada Mountains clade. These results corroborate previous inferences made with restriction enzyme-based approaches (Stone and Wolfe 2021; Stone et al. 2023). While the 10kb window ASTRAL-based species tree and the concatenated ML tree produced identical topologies, the CDS ASTRAL-based tree differed in the placement of one branch, placing *P. rupicola* from Cle Elum Lake in central Washington as sister to the remainder of the *P. rupicola* clade, instead of the sample from Castle Lake in northern California (Figure S.1). From this point forward we refer to the 10kb window ASTRAL tree as the species tree (Figure 1). Despite agreement among alternative species tree topologies and high bootstrap values, quartet support and concordance factors (gCFs and SCFs) were relatively low (Figure 1). This suggests conflicting phylogenetic signal among window trees (due to ILS and/or introgression) at many branches, especially for relationships deeper in the tree. While the overall level of discordance was relatively high, the most strongly supported internal branch (with the least discordance) is the branch placing *P. newberryi* and *P. cardwellii* as sister (Figure 1, IB2: quartet support = 50%, gCF = 18.6%, sCF = 58.8%), suggesting strongly that the two hummingbird-pollinated species (*P. newberryi* and *P. rupicola*) are not sister species.

**Figure 1.**
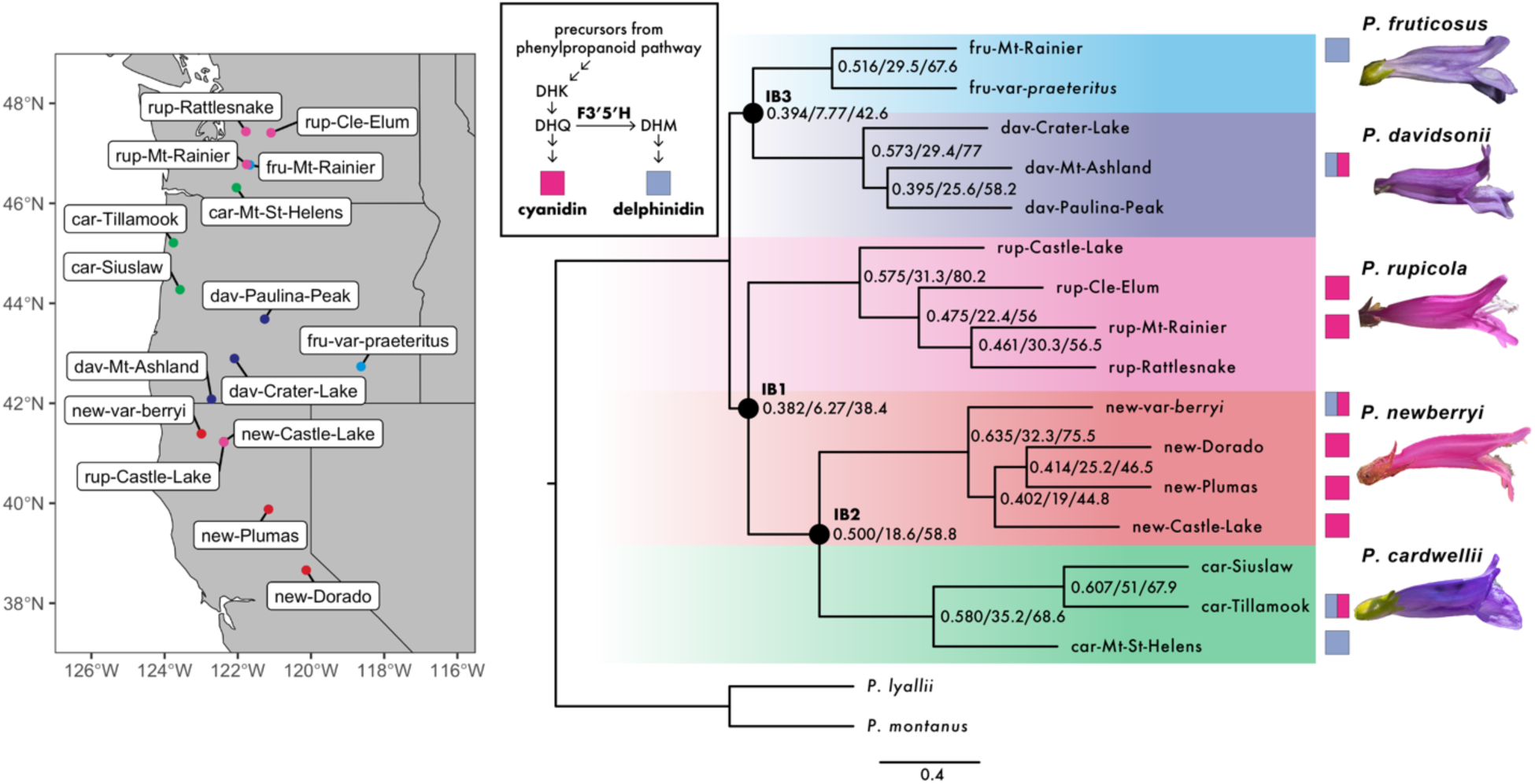
Sampling sites, species relationships, and anthocyanin pigmentation data for sampled *Dasanthera* taxa. Left: sampling localities for all ingroup samples included in this study. Right: 10kb genomic window ASTRAL species tree, where tips are individual samples as shown on the map. Numbers on nodes indicate ASTRAL quartet support / gene concordance factors / site concordance factors, respectively. All ASTRAL local posterior probabilities are 1. Black circles at select nodes denote focal internal branches. Magenta and blue-violet squares indicate floral anthocyanidins produced by each sample; pigment extraction was unsuccessful for samples without squares. Inset shows a simplified cartoon version of the anthocyanin biosynthesis pathway where Flavonoid 3’,5’-Hydroxylase (F3’5’H) functions to convert the precursor of cyanidin (dihydroquercetin; DHQ) into the precursor of delphinidin (dihydromyricetin; DHM). Arrows indicate enzymatic reactions.

Anthocyanin pigment data for the ten available samples confirm that the magenta flowers associated with bird pollination in *P. newberryi* and *P. rupicola* contain pink cyanidin-based anthocyanin pigments only, whereas bee-adapted species produce blue-delphinidin-based pigments, sometimes in combination with cyanidin (results summarized in Table S.3.). The only exception is the *P. newberryi* var. *berryi* individual that produced both cyanidin and delphinidin.

Flowers of this taxon are not nearly as apparently specialized to bird pollination: the anthers and stigma are inserted within the flower, and the corolla tube may appear blue-violet in color, rather than magenta. Overall, these results align with a previous study reporting a single *P. newberryi* sample produced cyanidin and a single *P. montanus* sample produced both cyanidin and delphinidin (Scogin and Freeman 1987).

Because the production of cyanidin is biochemically simpler than the production of delphinidin, *P. rupicola* and *P. newberryi* (with the exception of the var. *berryi* sample) have experienced a loss of delphinidin production, relative to the bee-pollinated species. Taking the species tree at face value, these results would suggest one of the following scenarios: (1) parallel losses of the ability to produce delphinidin (resulting in magenta flowers) in *P. newberryi* and *P. rupicola*, (2) a single loss of delphinidin production along the branch leading to *P. newberryi*, *P. cardwellii*, and *P. rupicola*, followed by at least one reversal in *P. cardwellii* (and additionally introgression into *P. newberryi* var. *berryi*, or a second reversal in this taxon), or (3) a single loss of delphinidin production, which is shared by *P. newberryi* and *P. rupicola* through either hemiplasy (ILS) or introgression.

### Coding sequence variation at F3′5′h suggests separate LOF de novo mutations are responsible for the repeated evolution of magenta flowers

Nucleotide sequence alignments of the *F3′5′h* gene sequence revealed that a diversity of LOF mutations to the coding sequence accompany loss of delphinidin production in *P. newberryii* and *P. rupicola*. We identified two distinct large deletions (∼500 bp) present in two *P. newberryi* samples from Eldorado and Plumas National Forests but absent in all other samples of *P. newberryi* and *P. rupicola* (Figure 2c). These deletions either partially or entirely remove the second exon, undoubtedly abolishing the activity of F3′5′H (Seitz et al. 2007). A third *P. newberryi* sample from Castle Lake shares a 4-bp insertion with the sample from Plumas which produces a premature stop codon (Figure 2c; Figure S.2). All three of these samples only produce cyanidin in their floral tissues (Table S.3). The *F3′5′h* sequence for *P. newberryi* var. *berryi* lacks inactivating mutations, consistent with observed delphinidin production in its floral sample. Two *P. rupicola* samples from Mt. Rainier and Rattlesnake Ridge shared a 4-bp insertion that results in premature stop codons. We were unable to obtain pigment data for the *P. rupicola* sample from Rattlesnake Ridge, but the sample from Mt. Rainier produces only cyanidin in floral tissues (Table S.3). The remaining two *P. rupicola* samples are less clear. One of these samples (from Cle Elum Lake) produces only cyanidin in flowers and is heterozygous for sites that cause amino acid substitutions in SRS1 and the hydroxylation activity site (Figure S.2) – these replacement mutations could reduce functionality of F3′5′H. The *P. rupicola* sample from Castle Lake has no suggestive functional mutations in *F3′5′h*, but we were unable to extract anthocyanidins due to poor tissue quality.

**Figure 2.**
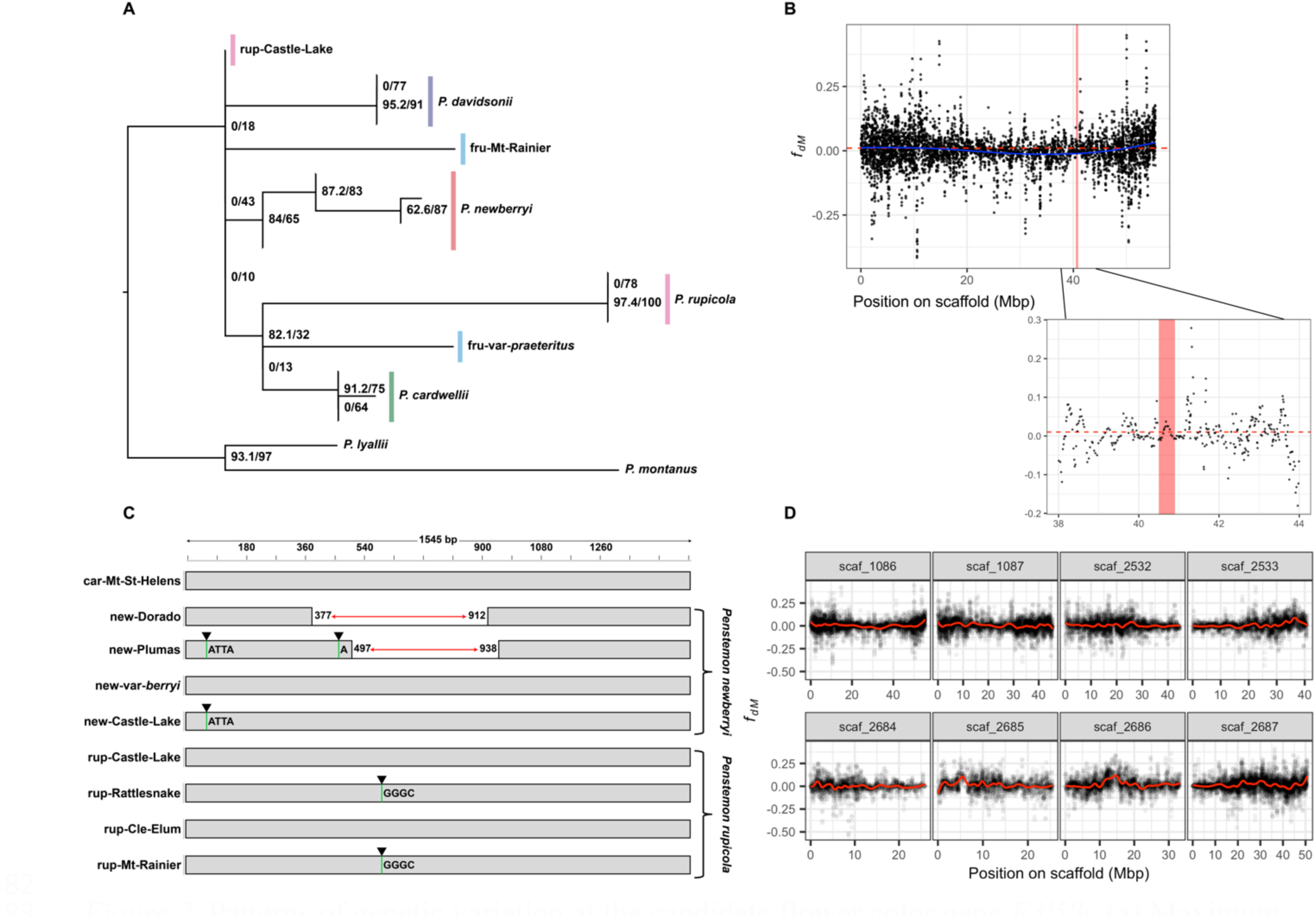
Patterns of genetic variation at the candidate flower color gene *F3’5’h*. (a) Maximum-likelihood gene tree estimated for *F3′5′h* gene sequence. Numbers at nodes signify the Shimodaira-Hasegawa approximate likelihood ratio test (SH-aLRT) support (%) / ultrafast bootstrap support (%). Branches with no SH-aLRT support are collapsed. (b) Values of the modified *f_d_* introgression statistic (*f_dM_*) for the triplet *P. cardwellii*-*P. newberryi*-*P. rupicola*, calculated in 500 SNP windows, sliding every 250 SNPs. Positive values are consistent with introgression between *P. newberryi* and *P. rupicola*, and negative values are consistent with introgression between *P. cardwellii* and *P. rupicola*. *f_dM_* is symmetrically distributed around zero under the null hypothesis of no introgression. The red bar indicates a 400kb window surrounding *F3′5′h*, the red dashed line indicates the scaffold-wide average *f_dM_* value. (c) Visualization of CDS for *F3′5′h* locus for a reference *P. cardwellii* (known functional copy) and all *P. newberryi* and *P. rupicola* samples. Red arrows represent deletions. Green lines with black arrows represent frameshift insertions. (d) Scaffold-by-scaffold values of *f_dM_*, with the same parameters as in panel B. Red lines are loess-smoothed *f_dM_* values.

Values of *f_dM_* assessing introgression between *P. newberryi* and *P. rupicola* surrounding the *F3′5′h* locus were near zero and were not significantly elevated above the scaffold-wide mean (one-tailed t-test: *p*-value = 0.18), suggesting *F3’5’h* is unlikely to have been shared between the two species through introgression (Figure 2b). This pattern is apparent regardless of the genomic window size used (Figure S.3). The maximum likelihood *F3′5′h* gene tree was discordant from the species tree, but not in a manner consistent with introgression of between *P. newberryi* and *P. rupicola* (Figure 2a). However, we note that many internal branches of the tree were quite poorly supported, likely due to low information content. In summary, we identified several distinct LOF mutations in *F3′5′h*. Because LOF mutations are difficult to reverse through gene repair, we find a scenario involving a single loss of delphinidin followed by reversals through gene repair (in *P. cardwellii* and *P. newberryi* var. *berryi*) to be highly unlikely. Further, we find no evidence for a history of introgression at the *F3′5′h* locus. Our data are consistent with parallel, independent LOF mutations as the mechanism behind the repeated evolution of magenta flowers in *Dasanthera*.

### Introgression is prevalent in Dasanthera, but does not suggest a history of gene flow between hummingbird-pollinated species

Our analysis of genome-wide introgression found that many taxa exhibit a history of introgression and admixture, consistent with previous inquiries into the group (Stone and Wolfe 2021; Stone et al. 2023). 64% of ABBA-BABA tests for all trios of samples in our dataset were significantly different from zero (after a Bonferroni correction at *α* = 0.05), providing strong support for widespread gene flow between species (Table S.4). The f-branch statistics corroborated these results and identified individual samples with elevated levels of introgression with another taxon, suggesting population-specific admixture that in some cases may be recent (Figure 3). Included among these results are (1) the *P. rupicola* sample from Cle Elum Lake, with introgression from *P. fruticosus*, (2) the *P. rupicola* sample from Castle Lake, with introgression from *P. newberryi*, and (3) the *P. davidsonii* sample from Crater Lake, with introgression from the clade containing *P. newberryi*, *P. cardwellii*, and *P. rupicola* (Figure 3). In each of these examples, the taxa in question are geographically close enough to exchange genes. As such, these may represent lineages that have experienced relatively recent bouts of introgression. Additionally, we identified significant signatures of introgression between *P. davidsonii* and both *P. newberryi* and *P. rupicola*, a result consistent with the presence of longstanding hybrid zones between these species.

**Figure 3.**
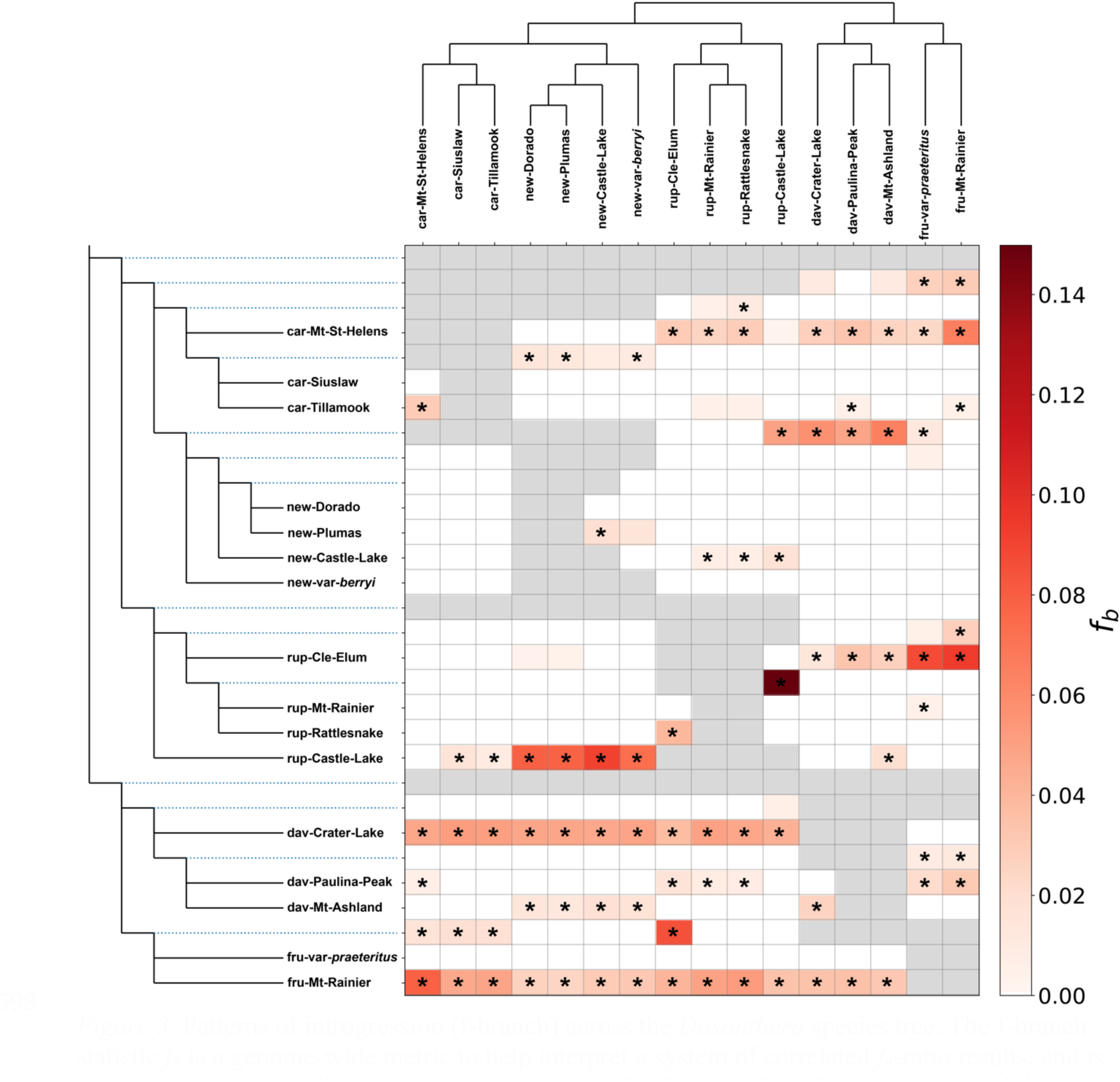
Patterns of introgression (f-branch) across the *Dasanthera* species tree. The f-branch statistic *f_b_* is a genome-wide metric to help interpret a system of correlated *f_4_*-ratio results, and is interpreted as excess allele (i.e., introgression) sharing between branches *b* (y-axis) and tips (x-axis) in the species tree. The color gradient represents the *f_b_* score; grey boxes represent tests that are inconsistent with the species tree topology (tip is descended from sister branch of focal branch *b*). Statistically significant results (Z-score > 3) are indicated by an asterisk.

While our *F3′5′h* sequence analyses suggest independent parallel evolution of magenta flowers in the two hummingbird-adapted species, the repeated evolution of other floral traits that make up the hummingbird syndrome (e.g., narrow floral tubes, increased nectar, elongated reproductive organs) might have evolved through adaptive introgression. In this scenario, we might expect to see a prevailing genomic signal of admixture between *P. newberryi* and *P. rupicola*. Contrary to this, our f-branch results find only limited support for introgression between *P. newberryi* and *P. rupicola*. Introgression between these species appears to be limited to samples collected together at the same site near Castle Lake in the Trinity Mountains of northern California. This region represents the northernmost and southernmost extent of the distributions of *P. newberryi* and *P. rupicola* respectively. Genome-wide estimates of the D statistic suggest a slightly positive but statistically insignificant signature of introgression between *P. newberryi* and *P. rupicola* (D = 0.003, *p* = 0.409; Table S.5). Moreover, *f_dM_* estimated in windows across the genome fail to identify any clear signatures of introgression between these species (Figure 2d). After removing the two samples from Castle Lake, genome-wide estimates of the D statistic suggest a significant but weak signature of introgression between *P. rupicola* and *P. cardwellii* (D = −0.037, *p* = 2.30e^−16^).

### Genome-wide patterns of genomic variation indicate selection against introgression

Genome-wide patterns of genetic divergence and discordance in *Dasanthera* are strongly shaped by local gene density (Figures S.4-S.7). We found that both genetic distance (*d_xy_*) and differentiation (*F_ST_*) between species are significantly positively associated with gene density (Table S.6), a pattern that is evident in both 10kb and 50kb windows (Figure S.8). Conversely, normalized Robinson-Foulds distance (a measure of topological discordance) of trees calculated from genomic windows relative to the species tree was negatively related to gene density, such that RF distance decreases (window trees become more similar to the species tree) as genic fraction increases (Figure 4). The topology weights for the main internal branches of the species tree were higher in genomic regions with a larger proportion of coding sequence (Figure 4), suggesting a higher fraction of taxa have data conforming to the species tree in gene dense regions. Moreover, topology weights for the internal branch supporting *P. newberryi* and *P. cardwellii* as sister taxa (IB2) were consistently higher across the genome compared to the topology weights for the other two internal branches. This pattern is consistent with metrics of branch support (quartet weights and concordance factors) that find stronger support overall for IB2 than for either of the other internal branches (Figure 4).

**Figure 4.**
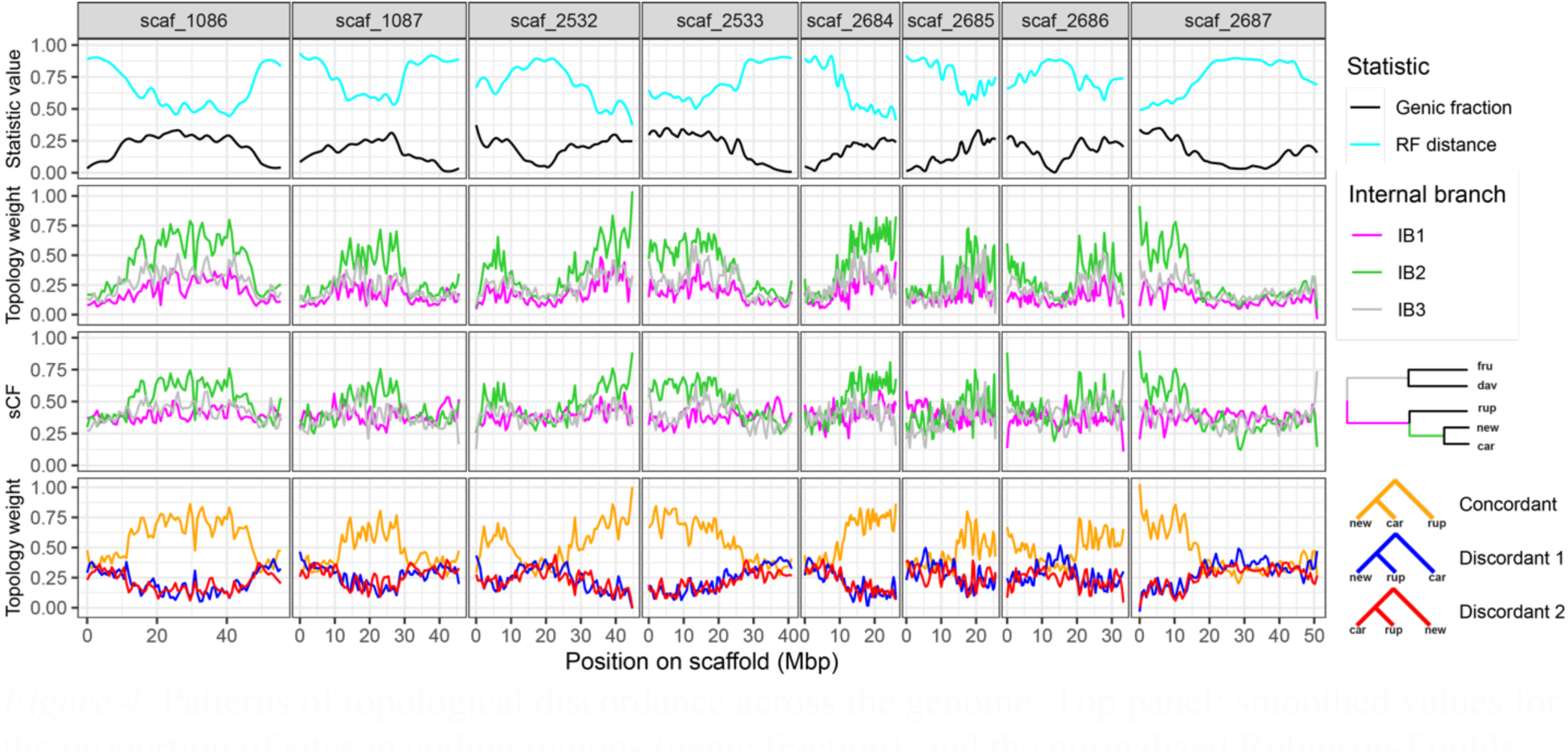
Patterns of topological discordance across the genome. Top panel: smoothed values for the proportion of sites in coding regions (genic fraction), and the normalized Robinson-Foulds distance between 10kb window trees and the species tree. Second panel: smoothed topology weights for 5-taxon window trees consistent with the three main internal branches of the species tree. Third panel: Smoothed site concordance factors (sCF) averaged over 100 quartets for 10kb window trees at the three main internal branches of the species tree. Bottom panel: Smoothed topology weights for the triplet: *P. newberryi* (*new*), *P. cardwellii* (*car*), *P. rupicola* (*rup*).

In general, we found that introgression is lower in more gene-dense regions of the genome (Figure S.9), consistent with selection against introgression. For five of the eight possible interspecific comparisons, there was a significant negative relationship between *f_dM_* and genic fraction. Curiously, this negative relationship was absent for all three comparisons including *P. fruticosus*: two comparisons showed no relationship, and one comparison showed a significantly positive relationship. Two species pairs are known to form hybrid zones in secondary contact: *P. davidsonii*-*P. newberryi* and *P. davidsonii*-*P. rupicola*. We found strong and significant negative relationships between *f_dM_* and genic fraction in both cases. This signal was stronger in *P. davidsonii*-*P. newberryi* (regression coefficient = −0.119; *R^2^* = 0.115) than in *P. davidsonii*-*P. rupicola* (regression coefficient = −0.065; *R^2^* = 0.054) and in both cases is consistent with selection against introgression. We additionally searched for genomic windows (10kb sliding intervals) exhibiting exceptionally high levels of genetic differentiation. We found a total of 70 outlier *d_xy_* windows that were present in both species pairs (shared outlier windows). These shared outlier windows were distributed across each of the eight largest scaffolds in the *P. davidsonii* reference genome (Figure S.10). Given their high degree of divergence in the face of hybridization, these regions may be particularly resistant to gene flow and may represent genomic loci underlying reproductive barriers in both hybrid zones.

## Discussion

### Independent loss-of-function mutations contribute to repeated adaptation in an adaptive radiation

Shifts from blue-violet flowers to magenta flowers in two *Dasanthera* species are most likely the result of independent LOF mutations to a gene in the anthocyanin biosynthesis pathway, *F3′5′h*. First, this gene is required for the production of delphinidin-based pigments and non-functional alleles are present in the magenta-flowered species that lack delphinidin pigments. This result strongly suggests shifts to magenta flowers involve LOF alleles at this gene. Second, distinct LOF alleles are found in *P. newberryi* and *P. rupicola*. Finally, we found no signal of introgression between the two magenta-flowered species near the *F3′5′h* gene, based on the genealogy and values of *f_dM_* surrounding the gene. These findings highlight the role of de novo mutation in the repeated evolution of adaptive phenotypes in an adaptive radiation of plants, which is in contrast to a growing body of research suggesting that parallel evolution in adaptive radiations is often derived from the adaptive introgression of pre-existing genetic variation (Colosimo et al. 2005; Schluter and Conte 2009; Berner and Salzburger 2015; Marques et al. 2019). It is worth considering that the type of mutation – in this case, one conferring loss-of-function – may strongly influence the frequency of de novo mutation in driving parallel phenotypic evolution. LOF mutations are an abundant source of variation in natural populations and may act as hotspots for repeated evolutionary transitions (Martin and Orgogozo 2013; Xu and Guo 2020). For example, LOF mutations are particularly prevalent in the evolution of plant-pollinator interactions; the loss of scent compounds, associated with transitions to hummingbird pollination and self-pollination, has repeatedly and independently evolved in several plant groups, including in *Petunia* (Segatto et al. 2014; Amrad et al. 2016), *Capsella* (Sas et al. 2016), and monkeyflowers (Peng et al. 2017; Liang et al. 2023). While there are many examples of LOF mutations conferring adaptive benefit, their occurrence is often limited to genes with few pleiotropic effects, or to regulatory elements that affect gene expression only in specific tissues (Xu and Guo 2020). Although transcription factors in the flavonoid pigment pathway have been shown to generally have higher rates of molecular evolution than the structural genes (like *F3′5′h*) they regulate (Wheeler et al. 2022), LOF mutations to *F3’5’h* seem to incur minimal pleiotropic effects, perhaps explaining their apparent preponderance. *F3’5’h* functions to produce tri-hydroxylated flavonoid compounds including delphinidin. Yet tri-hydroxylated flavonoids seem to be restricted to floral tissue – many species that have delphinidin-producing flowers produce di-hydroxylated flavonoids (products of the upstream gene *F3′h*) in vegetative tissues (Wessinger and Rausher 2012). Furthermore, several plant species have evolved red floral pigments through LOF mutations in *F3′5′h* (Smith and Rausher 2011; Ishiguro et al. 2012) including other species of *Penstemon* (Wessinger and Rausher 2015), and many other angiosperm taxa lack this gene entirely (Wessinger and Rausher 2012), suggesting the defunctionalization of *F3′5′h* can underlie adaptation without deleterious side-effects. While transitions to magenta flowers in *P. newberryi* and *P. rupicola* appear to involve parallel de novo mutations, this does not preclude a role for adaptive introgression fueling other aspects of the hummingbird floral syndrome. However, we found no clear evidence of gene flow between *P. newberryi* and *P. rupicola* genome-wide or near the *F3′5′h* locus. While these results do not rule out the potential for adaptive introgression of other floral traits associated with hummingbird pollination syndrome (e.g., exserted stigma and anthers), at this time we have no evidence to support this hypothesis.

It is also worth considering that the degree of color shift we observe in *Dasanthera* is not as severe as in other *Penstemon* taxa that have transitioned to hummingbird pollination. Most other hummingbird-pollinated *Penstemon* species produce pelargonidin-based anthocyanins (Scogin and Freeman 1987; Wessinger and Rausher 2015), whereas the hummingbird-adapted *Dasanthera* species produce the more-hydroxylated cyanidin. This difference is observable by the naked eye, such that flowers with cyanidin are pink-magenta, while flowers with pelargonidin are red. It is possible that the absence of pelargonidin in *Dasanthera* flowers reflects a more generalized floral phenotype that is thus less specialized to hummingbird pollination. Although hummingbirds are frequent visitors to *P. newberryi*, they are only present at low and mid-elevations throughout their range; a diverse community of potential pollinators visit both *P. newberryi* and *P. davidsonii*, which suggests a somewhat generalist phenotype in *P. newberryi* and explains why hybridization is so common between these species (Kimball 2008).

And, while *P. newberryi* has many characteristics of hummingbird specialization, other features, including small quantities of nectar (Scogin and Freeman 1987; personal observation) and the aforementioned magenta corolla suggest it has not fully transitioned to a specialized hummingbird-pollination phenotype. The same may be true for *P. rupicola*, where (to the authors’ knowledge) the only pollinator observation study found that while hummingbirds do visit *P. rupicola* flowers, the most common floral visitors to *P. rupicola* were pollen-collecting halictid bees, which are unlikely to be effective pollinators (Datwyler 2001). Conversely to *P. newberryi*, however, *P. rupicola* produces large quantities of nectar (personal observation), consistent with specialization to hummingbird pollination. In both cases, the apparent lack of complete pollinator specialization likely plays a large role in facilitating interspecific gene flow, an apparently common phenomenon in *Dasanthera*.

*Patterns of discordance and introgression are consistent with selection driving divergence* We found widespread signatures of introgression, corroborating previous inquiries into hybridization dynamics in *Dasanthera* (Stone and Wolfe 2021; Stone et al. 2023). However, patterns of introgression and phylogenetic discordance were heterogeneous across the genome. The genomic landscape of discordance and introgression are consistent with selection driving species divergence and opposing introgression in the *Dasanthera* radiation. Trees inferred from genomic windows tended to be more similar to the species tree topology in regions of higher gene density (Figure 4). This relationship between discordance and gene content is consistent with stronger purifying selection acting in gene dense regions: purifying selection acts to reduce effective population size hence reducing discordance caused by ILS, and acts to oppose introgression, a further source of discordance. Moreover, gene-dense regions exhibited greater differentiation and reduced signals of introgression, consistent with reduced gene flow in gene-rich regions. We expect that levels of differentiation and introgression are also shaped by variation in local recombination rate, a near-universal pattern in population genetic datasets (Begun and Aquadro 1992; Cruickshank and Hahn 2014; Martin and Jiggins 2017; Wolf and Ellegren 2017). Specifically, high recombination regions should experience reduced genetic differentiation and tolerate higher levels of introgression. Unfortunately, we currently lack a recombination map for the *P. davidsonii* reference genome. In a more distantly related *Penstemon* species (*P. barbatus*), gene-dense regions tend to have higher recombination rates, resulting in an overall negative relationship between population differentiation and recombination rate (Wessinger et al. 2023).

The two hummingbird-adapted species in our study both hybridize with the bee-adapted species *P. davidsonii* in the Cascades and Sierra Nevada Mountains. These sets of species pairs provide an opportunity to examine whether similar genomic barriers to gene flow accompany parallel divergence in floral syndrome. For each species pair, we found strongly differentiated genomic regions (Figure S.10). Such localized genomic regions of elevated divergence are expected when there is strong divergent selection in the face of gene flow (Charlesworth et al. 1997; Feder and Nosil 2010). These genomic loci appear resistant to introgression and may potentially be involved in local adaptation to divergent habitats or reproductive isolation. One caveat is that these outlier loci do not truly represent independent regions of elevated divergence, because they include the same samples of *P. davidsonii* in both contrasts. Future genomic and phenotypic analyses of these replicate hybrid zones will be better suited to determine whether these genomic regions represent parallel loci of elevated divergence, and will begin to address whether patterns of phenotypic variation and selection within hybrid zones show similar and/or predictable patterns.

## Conclusion

*Penstemon* subgenus *Dasanthera* represents a microcosm adaptive radiation, with its nine species exhibiting a diversity of ecological niches and floral pollination strategies. Species boundaries remain porous in *Dasanthera*, allowing for interspecific hybridization and highlighting the potential for the adaptive introgression of ecologically important traits. In this study we focused on the source of genetic variation for repeated origins of a novel flower color in hummingbird-pollinated species, as we currently lack information on the genetic basis for other ecological traits that have diversified in the group. Although our data suggests parallel de novo mutations underlie repeated flower color evolution, it is plausible that adaptive introgression has fueled other aspects of the radiation because evidence for introgression is prevalent in *Dasanthera*. We suspect that both parallel de novo mutations and introgression have contributed to diversification, as inferred in other rapidly radiating clades.

## Acknowledgements

We thank members of the Wessinger lab for discussions on the project and the USFS and NPS for sampling permits. We thank Kate Ostevik and Mark Rausher for access to the *P. davidsonii* genome prior to its publication. The Division of Information Technology at the University of South Carolina provided access to computing resources on the Hyperion HPC cluster. We thank Stacey Smith and an anonymous reviewer for helpful feedback on the manuscript. This work was funded by NSF IOS-2209128 (to B.W.S.), NSF DEB-2052904 (to C.A.W), and NIH NIGMS R35GM142636 (to C.A.W.). Scripts used for this study are available at https://github.com/benstemon/MBE-23-0936.

## Data Availability Statement

Raw sequencing data have been deposited into the NCBI Sequence Read Archive under the accession number PRJNA1057825. Other data underlying this article, including processed .vcf files, are available on figshare under doi: 10.6084/m9.figshare.24480499.

