## Supplemental Figures and Tables for "Ecological diversification in an adaptive radiation of plants: the role of de novo mutation and introgression"

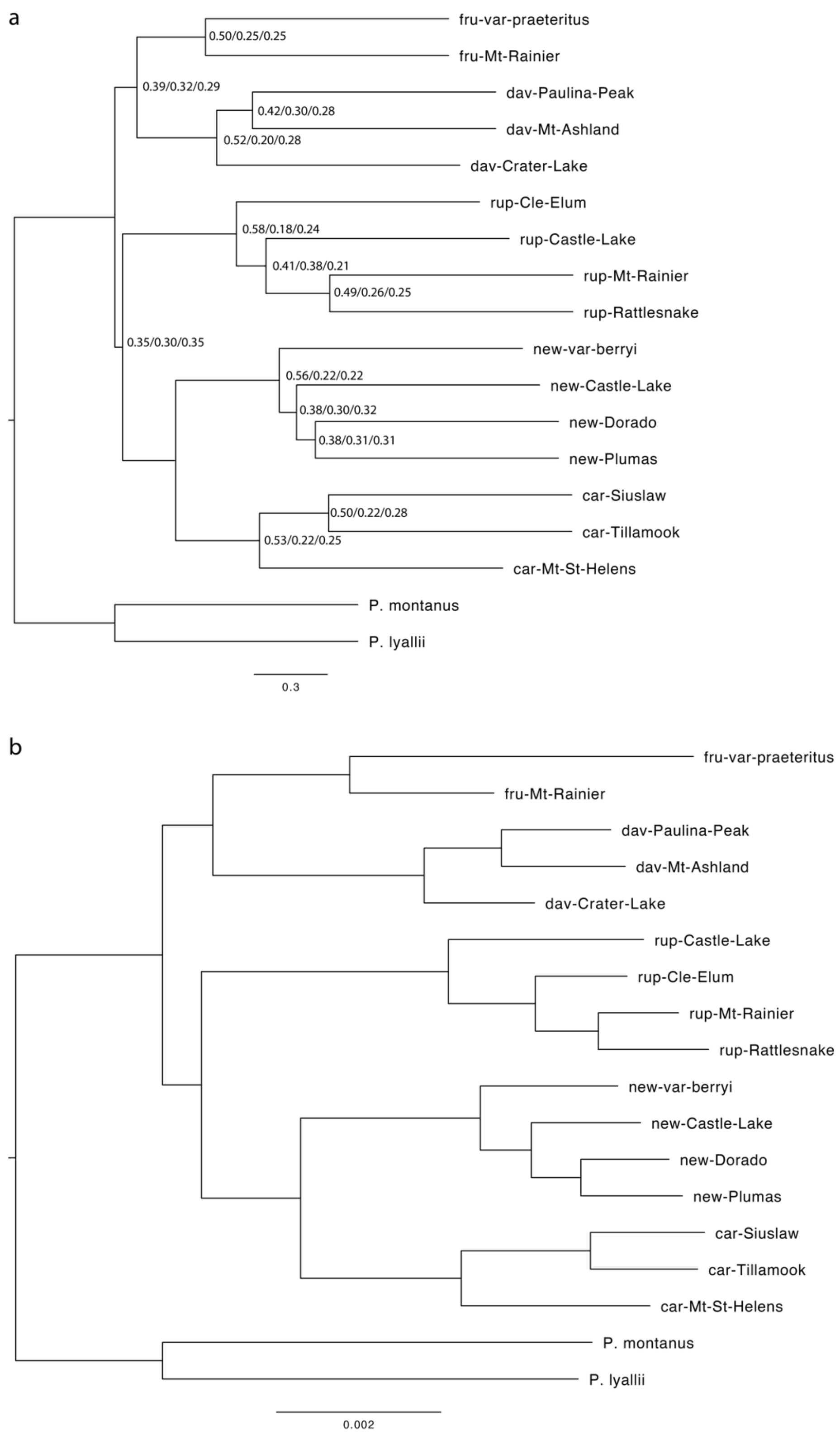


**Figure S.1.** Alternative species trees. (a) ASTRAL CDS tree. Labels at nodes indicate ASTRAL quartet scores for q1, q2, and q3, respectively. All ASTRAL local posterior probabilities are 1. (b) Concatenated MLE tree. All UFBoot values are 100%.

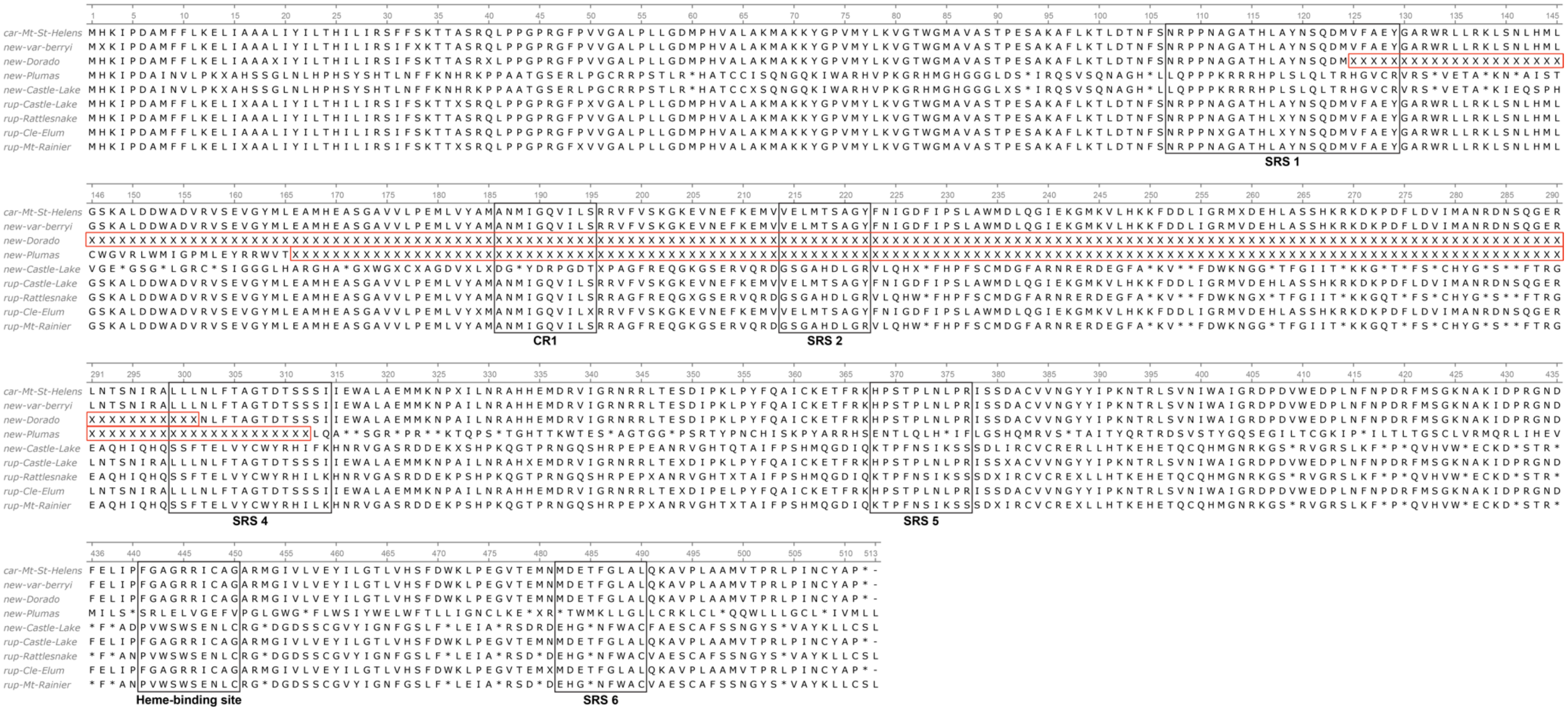


**Figure S.2.** Protein alignment of *F3'5'h* for a reference *P. cardwellii* (known functional copy) and all samples of *P. newberryi* and *P. rupicola*. Black boxes outline substrate recognition sites (SRS), the heme-binding domain, and the conserved hydroxylation activity site (CR1). We were unable to definitively identify the position of SRS3 through sequence homology. Red boxes outline large deletions in two of the *P. newberryi* samples.

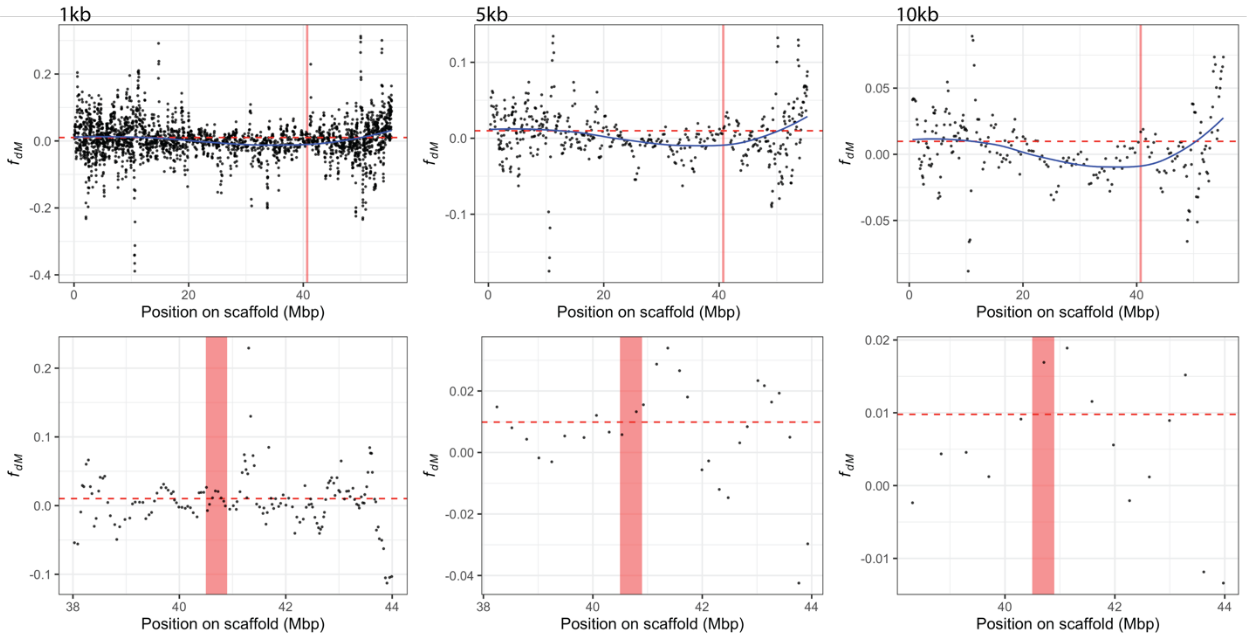


**Figure S.3.** Values of $f_{dM}$ in non-overlapping windows surrounding the *F3'5'h* locus. Window sizes are 1,000 SNPs, sliding every 250 SNPs (left), 5,000 SNPs, sliding every 1,250 SNPs, (center), and 10,000 SNPs, sliding every 2,500 SNPs (right).

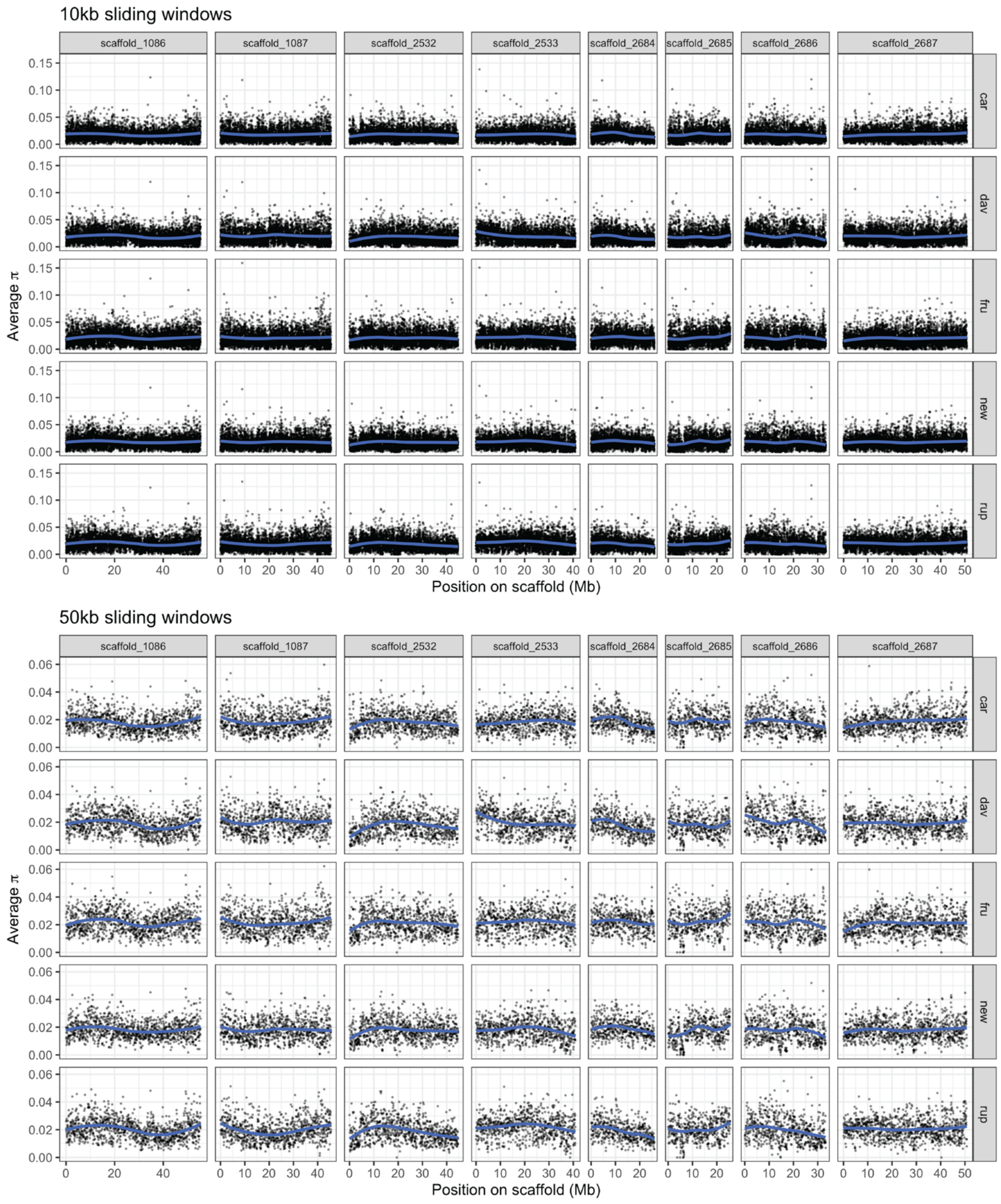


**Figure S.4.** Values of π across the eight main scaffolds of the *P. davidsonii* reference genome in 10kb (top) and 50kb (bottom) windows.

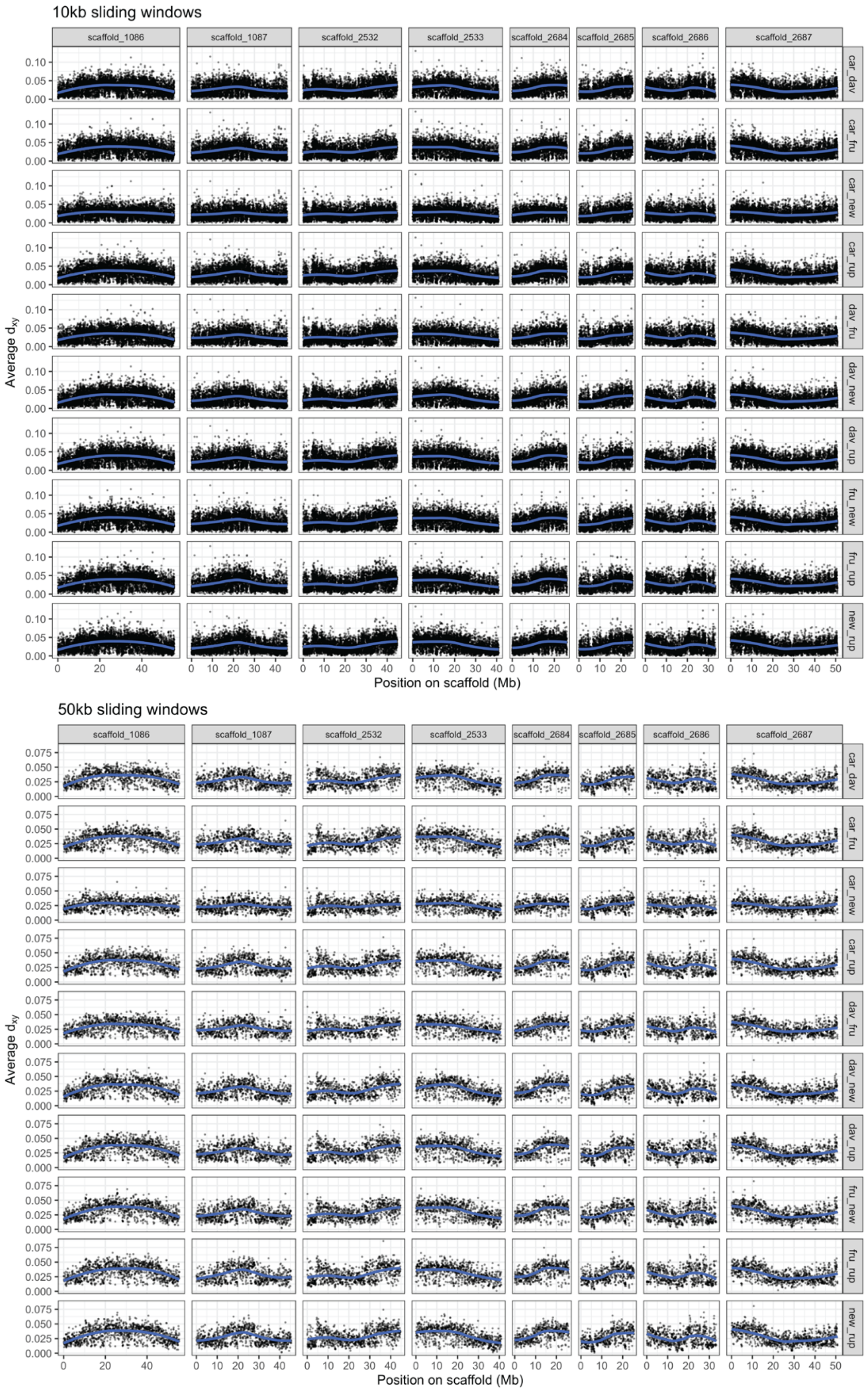


**Figure S.5.** Values of $d_{xy}$ across the eight main scaffolds of the *P. davidsonii* reference genome in 10kb (top) and 50kb (bottom) windows.

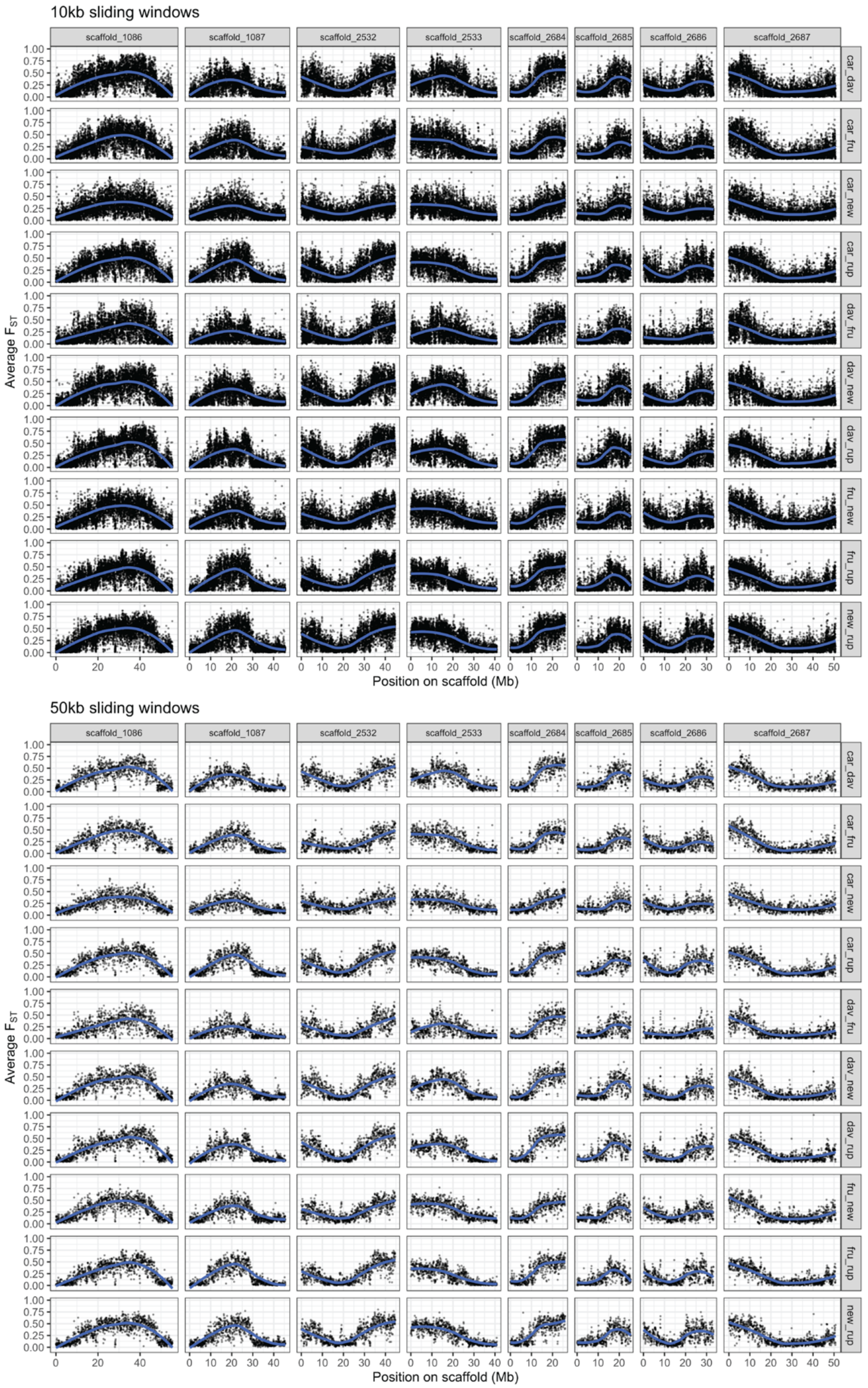


**Figure S.6.** Values of $F_{ST}$ across the eight main scaffolds of the *P. davidsonii* reference genome in 10kb (top) and 50kb (bottom) windows.

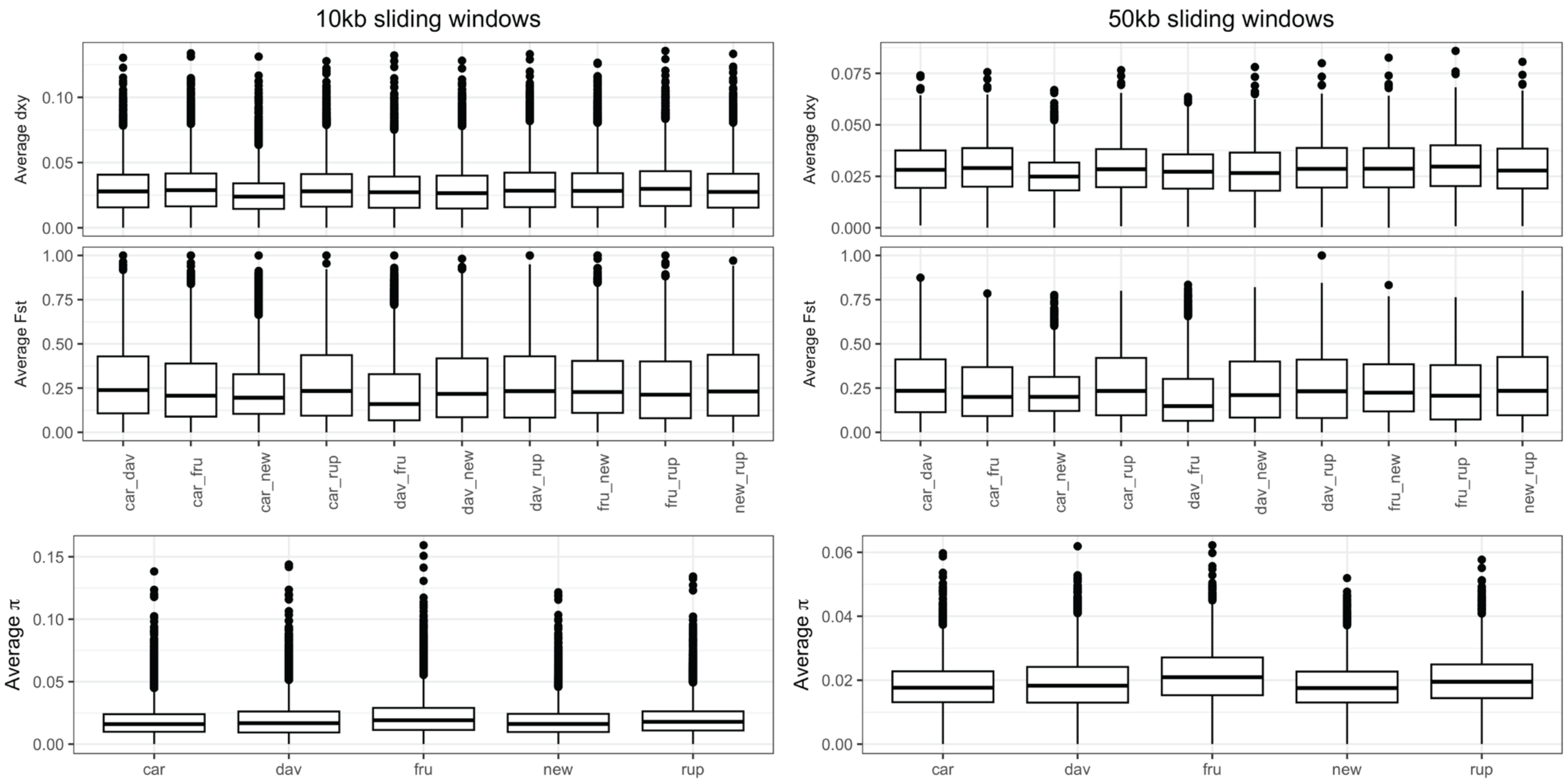


**Figure S.7.** Genome-wide values of $d_{xy}$, $F_{ST}$, and π in 10kb (left) and 50kb (right) windows.

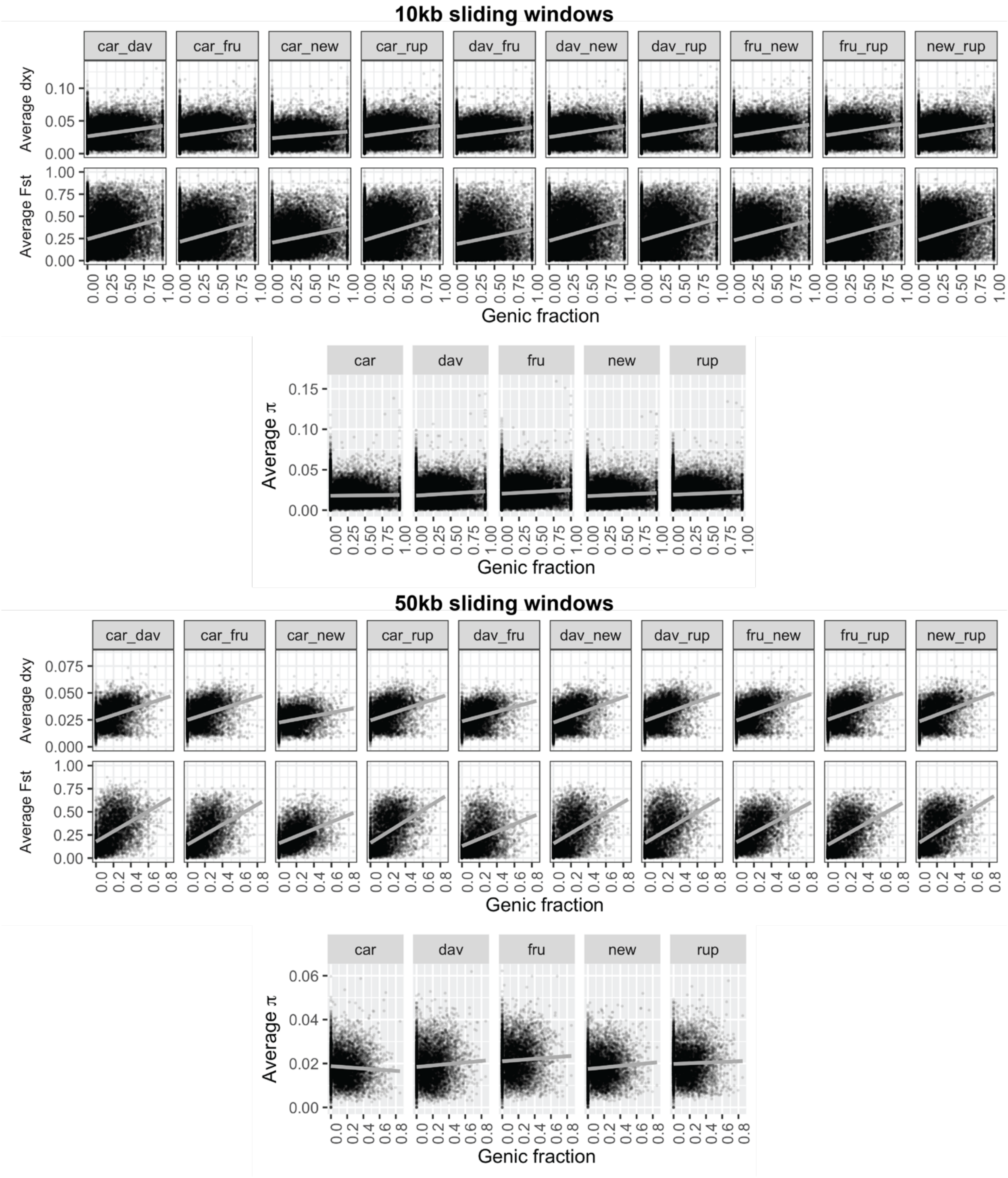


**Figure S.8.** Linear regressions between genic fraction and diversity and divergence metrics ($d_{xy}$, $F_{ST}$, and π) in 10kb (top) and 50kb (bottom) windows.

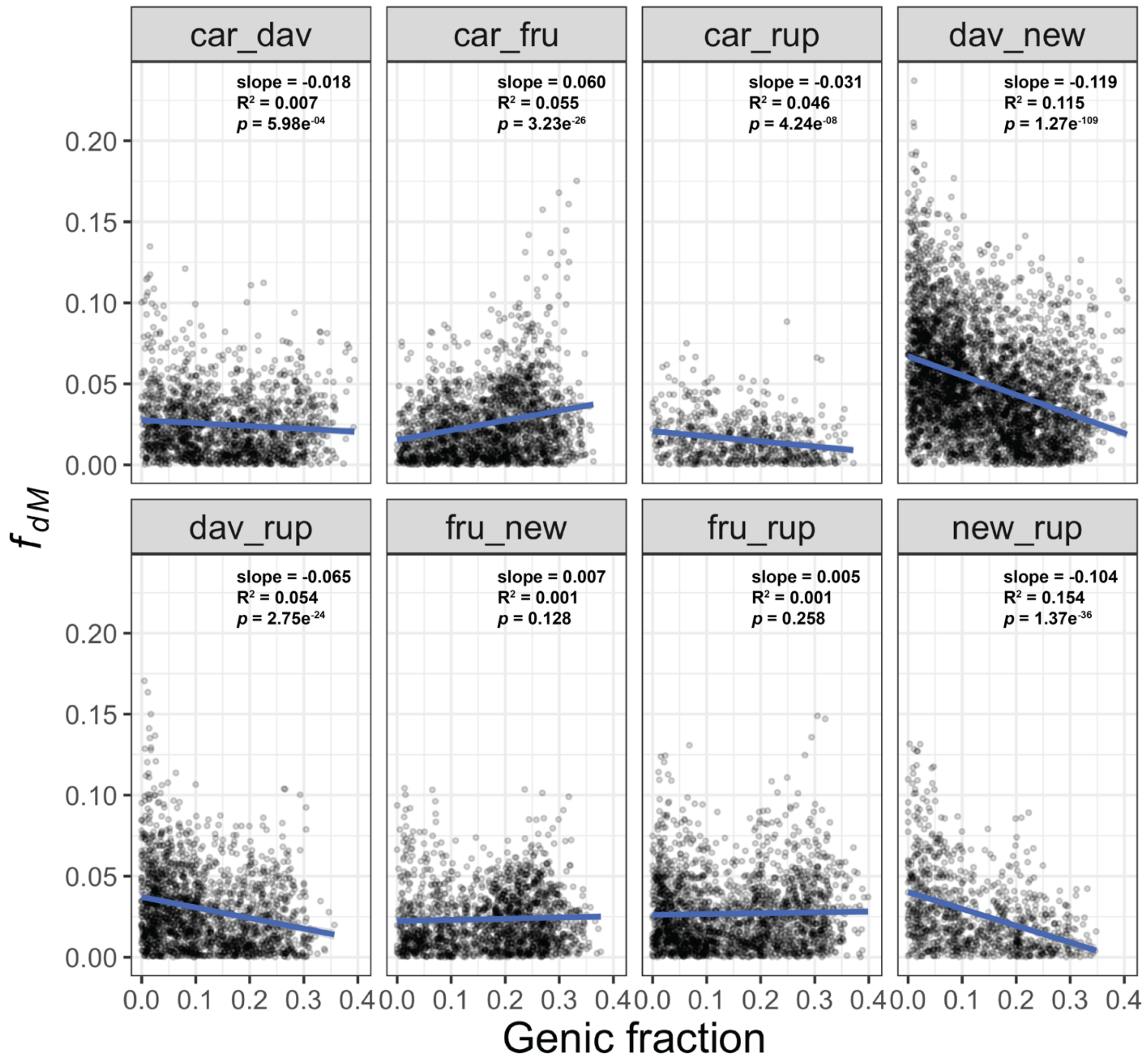


**Figure S.9.** Linear regressions between genic fraction and introgression ($f_{dM}$). $f_{dM}$ was calculated in 10,000 SNP windows, sliding every 2,500 SNPs.

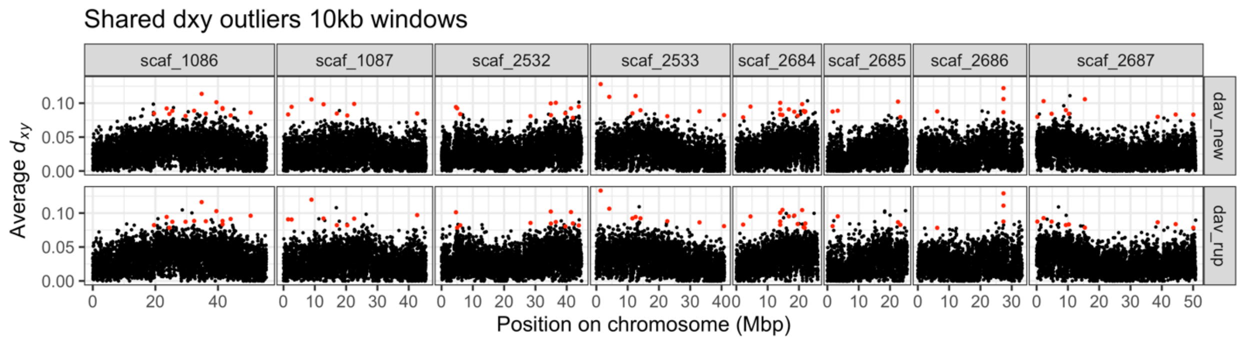


**Figure S.10.** $d_{xy}$ in sliding 10kb windows in two species pairs known to form hybrid zones (*P. davidsonii-P. newberryi* and *P. davidsonii-P. rupicola*). Outlier $d_{xy}$ values (Z-score >3) in the same windows for both species pairs are highlighted in red.

*Table S.1.* Sampling information

| Species | Identifier | Population ID | Latitude | Longitude |
|---|---|---|---|---|
| *Penstemon rupicola* | Castle Lake | 86-8 | 41.223113 | -122.379630 |
| | Rattlesnake | 98-4 | 47.436029 | -121.778519 |
| | Mt. Rainier | 105-2 | 46.778033 | -121.760309 |
| | Cle Elum | 101-1 | 47.411009 | -121.092183 |
| *Penstemon davidsonii* | Mt. Ashland | 87-7 | 42.081020 | -122.718583 |
| | Crater Lake | 116-7 | 42.895602 | -122.092252 |
| | Paulina Peak | 117-2 | 43.684535 | -121.265441 |
| *Penstemon newberryi* | Dorado | 75-9 | 38.661708 | -120.130671 |
| | Plumas | 80-5 | 39.880031 | -121.161161 |
| | var. *berryi* | 84-4 | 41.389587 | -122.993862 |
| | Castle Lake | 85-9 | 41.231111 | -122.382305 |
| *Penstemon cardwellii* | Mt. St. Helens | 28-5 | 46.313721 | -122.036324 |
| | Siuslaw | 91-6 | 44.274483 | -123.579297 |
| | Tillamook | 92-7 | 45.210899 | -123.758863 |
| *Penstemon fruticosus* | Mt. Rainier | 106-1 | 46.768161 | -121.674510 |
| | var. *praeteritus* | 118-1 | 42.735390 | -118.633691 |
| *Penstemon lyallii* | Kaniksu | 44-9 | 48.371445 | -116.156739 |
| *Penstemon montanus* | Challis | 61-7 | 44.511154 | -114.374046 |

*Table S.2.* Read depth and coverage information.

| | | | Scaffolds | **1086** | **2687** | **1087** | **2532** | **2533** | **2686** | **2684** | **2685** | 1085 | 2151 | 2531 | 2446 | | |
|---|---|---|---|---|---|---|---|---|---|---|---|---|---|---|---|---|---|
| Species | Sample | reads mapped and paired | mean genome-wide depth | 6E+07 | 5E+07 | 5E+07 | 4E+07 | 4E+07 | 3E+07 | 3E+07 | 3E+07 | 1E+07 | 8E+06 | 7E+06 | 5E+06 | Total bp | Total coverage |
| *Penstemon rupicola* | rup-Castle-Lake | 26629396 | 11.8227 | 77.97% | 73.11% | 72.59% | 74% | 76.15% | 71.90% | 76.89% | 71.23% | 68.58% | 70.92% | 60.71% | 64.40% | 260102573 | 0.737 |
| | rup-Rattlesnake | 32852064 | 14.0107 | 77.59% | 71.70% | 72.79% | 74.55% | 74.69% | 70.36% | 76.98% | 68.43% | 68.18% | 72.45% | 53.78% | 68.60% | 257684226 | 0.730 |
| | rup-Mt-Rainier | 24422232 | 11.1638 | 77.89% | 71.17% | 71.71% | 73.31% | 74.72% | 70.30% | 76.71% | 68.05% | 68.18% | 68.89% | 57.75% | 67.24% | 256275894 | 0.726 |
| | rup-Cle-Elum | 26171512 | 11.5766 | 78.36% | 73.43% | 71.91% | 75.00% | 76.17% | 70.33% | 76.50% | 68.79% | 67.67% | 71.33% | 52.55% | 64.14% | 258900747 | 0.734 |
| *Penstemon davidsonii* | dav-Mt-Ashland | 27095921 | 11.8077 | 81.61% | 74.60% | 73.67% | 76.73% | 78.28% | 75.14% | 80.83% | 71.97% | 70.01% | 71.16% | 58.86% | 74.50% | 268470978 | 0.761 |
| | dav-Crater-Lake | 29184105 | 12.0241 | 81.39% | 75.15% | 74.09% | 76.72% | 76.87% | 72.41% | 80.67% | 71.86% | 70.37% | 72.43% | 56.92% | 72.82% | 267182235 | 0.757 |
| | dav-Paulina-Peak | 31564339 | 13.1641 | 81.71% | 75.01% | 73.93% | 76.37% | 77.35% | 75.46% | 80.42% | 71.87% | 71.61% | 69.99% | 58.10% | 74.93% | 268339775 | 0.760 |
| *Penstemon newberryi* | new-Dorado | 40276056 | 13.8701 | 81.33% | 76.65% | 77.14% | 76.90% | 79.13% | 75.47% | 80.06% | 76.51% | 72.38% | 74.76% | 67.81% | 68.57% | 273259508 | 0.774 |
| | new-Plumas | 36370992 | 15.2172 | 80.53% | 75.92% | 76.74% | 76.40% | 78.85% | 73.32% | 79.72% | 75.46% | 70.27% | 74.60% | 68.12% | 68.86% | 270644978 | 0.767 |
| | new-var-*berryi* | 29458522 | 12.2794 | 79.83% | 74.39% | 75.24% | 75.54% | 77.15% | 74.28% | 78.91% | 73.30% | 67.96% | 70.46% | 68.11% | 67.33% | 266618910 | 0.756 |
| | new-Castle-Lake | 24241868 | 10.8715 | 79.09% | 73.20% | 74.44% | 74.18% | 76.43% | 73.33% | 77.84% | 72.49% | 69.16% | 70.61% | 67.71% | 70.41% | 263807606 | 0.748 |
| *Penstemon cardwellii* | car-Mt-St-Helens | 36858317 | 16.006 | 78.78% | 72.62% | 73.08% | 74.79% | 74.86% | 71.02% | 78.11% | 68.85% | 67.87% | 71.91% | 58.15% | 65.45% | 259795581 | 0.736 |
| | car-Siuslaw | 28739053 | 13.0176 | 76.75% | 70.13% | 70.98% | 72.95% | 74.65% | 68.79% | 76.37% | 67.51% | 64.66% | 69.54% | 55.19% | 59.94% | 252987272 | 0.717 |
| | car-Tillamook | 32770333 | 13.5502 | 77.32% | 70.70% | 71.11% | 72.61% | 74.83% | 68.85% | 76.14% | 68.68% | 64.68% | 69.17% | 54.63% | 60.62% | 253803365 | 0.719 |
| *Penstemon fruticosus* | fru-Mt-Rainier | 21818078 | 9.9509 | 77.37% | 70.47% | 70.00% | 73.38% | 73.30% | 70.13% | 76.55% | 66.27% | 65.62% | 65.80% | 53.05% | 64.85% | 252795720 | 0.716 |
| | fru-var-*praeteritus* | 42247127 | 16.3052 | 77.71% | 71.17% | 71% | 73.33% | 73.58% | 70.27% | 76.89% | 66.96% | 67.76% | 69.03% | 51.65% | 69.97% | 254698494 | 0.722 |
| *Penstemon lyallii* | *P. lyallii* | 28008540 | 12.9174 | 74.10% | 66.74% | 64.43% | 69.32% | 69.11% | 64.44% | 73.37% | 61.40% | 60.49% | 63.77% | 37.92% | 57.22% | 236938120 | 0.671 |
| *Penstemon montanus* | *P. montanus* | 28564984 | 12.2487 | 72.85% | 64.82% | 64% | 67.48% | 69% | 63.18% | 71.48% | 61.70% | 59.95% | 60.07% | 41.67% | 56.54% | 233235972 | 0.661 |
| | **Total** | 547273439 | - | - | - | - | - | - | - | - | - | - | - | - | - | - | - |
| | **Average** | 30404080 | 12.878 | 0.785 | 0.723 | 0.721 | 0.741 | 0.753 | 0.711 | 0.775 | 0.695 | 0.675 | 0.698 | 0.568 | 0.665 | 258641220 | 0.733 |
| | **SD** | 5598161.809 | 1.729 | 0.024 | 0.031 | 0.035 | 0.025 | 0.028 | 0.034 | 0.025 | 0.041 | 0.034 | 0.036 | 0.083 | 0.054 | 10691775 | 0.030 |

After quality control, filtering, and mapping to the *P. davidsonii* reference genome, our whole-genome resequencing efforts produced 547.3 million paired-end reads, for an average of 30.4 ± 5.6 million reads per sample. Genome-wide depth of coverage was fairly consistent across samples (mean: 12.9x ±1.7x). The average per-scaffold coverage was > 70% for most of the eight largest scaffolds, while average coverage for the smaller four scaffolds was slightly reduced (56.8%-69.8%). Genome-wide coverage for most samples was > 70% (mean: 73.3% ±3.0%), with slightly lower coverage for the two outgroup species *P. montanus* and *P. lyallii* (66.1% and 67.1%, respectively).

*Table S.3.* Anthocyanin extractions from floral tissues.

| Species | Sample | Delphinidin | Cyanidin | Pelargonidin |
|---|---|---|---|---|
| *Penstemon rupicola* | rup-Castle-Lake | | | |
| | rup-Rattlesnake | | | |
| | rup-Mt-Rainier | - | + | - |
| | rup-Cle-Elum | - | + | - |
| *Penstemon davidsonii* | dav-Mt-Ashland | + | + | - |
| | dav-Crater-Lake | | | |
| | dav-Paulina-Peak | | | |
| *Penstemon newberryi* | new-Dorado | - | + | - |
| | new-Plumas | - | + | - |
| | new-var-*berryi* | + | + | - |
| | new-Castle-Lake | - | + | - |
| *Penstemon cardwellii* | car-Mt-St-Helens | + | - | - |
| | car-Siuslaw | | | |
| | car-Tillamook | + | + | - |
| *Penstemon fruticosus* | fru-Mt-Rainier | + | - | - |
| | fru-var-*praeteritus* | | | |
| *Penstemon lyallii* | *P. lyallii* | | | |
| *Penstemon montanus* | *P. montanus* | | | |

Colored boxes with a "+" symbol indicate the corresponding anthocyanidin was present in the sample. Black boxes indicate samples for which no anthocyanidins were successfully extracted.

*Table S.4.* Results of all possible ABBA-BABA tests.

| # of tests | Bonf. Corr | # significant | % significant |
|---|---|---|---|
| 550 | 9.0909E-05 | 352 | 0.64 |

Tests are not included in the calculation if P1, P2, and P3 all represent the same species.

*Table S.5.* ABBA-BABA tests for the triplet: car-new-rup

| **All samples included** | | | | | | | | |
|---|---|---|---|---|---|---|---|---|
| **P1** | **P2** | **P3** | **Dstatistic** | **Z-score** | **p-value** | **BBAA** | **ABBA** | **BABA** |
| car | new | rup | 0.00329235 | 0.825638 | 0.40901 | 527300 | 341400 | 339159 |

| **Castle Lake samples removed** | | | | | | | | |
|---|---|---|---|---|---|---|---|---|
| **P1** | **P2** | **P3** | **Dstatistic** | **Z-score** | **p-value** | **BBAA** | **ABBA** | **BABA** |
| car | new | rup | -0.0367873 | 12.5607 | **2.30E-16** | 531508 | 318122 | 342422 |

Top: All samples of *P. newberryi* , *P. cardwellii* , and *P. rupicola* were included. Bottom: The *P. newberryi* and *P. rupicola* samples from Castle Lake were removed.

*Table S.6.* Regression coefficients between gene density and genetic distance/divergence

| 50kb windows | | | | 10kb windows | | | |
|---|---|---|---|---|---|---|---|
| ***dxy*** | | | | | | | |
| species | coeff | r2 | pval | species | coeff | r2 | pval |
| car_dav | 0.02679961 | 0.14119495 | 2.76E-212 | car_dav | 0.01609035 | 0.06275877 | 0 |
| car_fru | 0.02636156 | 0.13185197 | 1.86E-192 | car_fru | 0.01584807 | 0.05928468 | 0 |
| car_new | 0.01537699 | 0.06985337 | 8.41E-103 | car_new | 0.01006378 | 0.03328736 | 1.60E-228 |
| car_rup | 0.0269757 | 0.13807176 | 3.57E-206 | car_rup | 0.01577318 | 0.05866475 | 0 |
| dav_fru | 0.02228994 | 0.1116958 | 6.06E-158 | dav_fru | 0.01467893 | 0.05656848 | 0 |
| dav_new | 0.02946169 | 0.16153785 | 2.17E-244 | dav_new | 0.01740973 | 0.07200574 | 0 |
| dav_rup | 0.02941799 | 0.15245021 | 2.33E-225 | dav_rup | 0.0176387 | 0.06949725 | 0 |
| fru_new | 0.02872283 | 0.1493096 | 9.15E-227 | fru_new | 0.01747815 | 0.0691676 | 0 |
| fru_rup | 0.02864688 | 0.13858072 | 2.00E-198 | fru_rup | 0.01722718 | 0.06453764 | 0 |
| new_rup | 0.03106926 | 0.16593077 | 3.25E-254 | new_rup | 0.01780928 | 0.07051457 | 0 |
| ***FST*** | | | | | | | |
| species | coeff | r2 | pval | species | coeff | r2 | pval |
| car_dav | 0.55050083 | 0.22293825 | 0 | car_dav | 0.23727683 | 0.08361573 | 0 |
| car_fru | 0.54103045 | 0.24688627 | 0 | car_fru | 0.23683177 | 0.09259425 | 0 |
| car_new | 0.38439291 | 0.20907734 | 0 | car_new | 0.16989382 | 0.06776272 | 0 |
| car_rup | 0.58889745 | 0.24979735 | 0 | car_rup | 0.25661709 | 0.09767121 | 0 |
| dav_fru | 0.39585108 | 0.14393333 | 1.35E-206 | dav_fru | 0.17170201 | 0.05242635 | 4.94E-324 |
| dav_new | 0.56023971 | 0.21803758 | 0 | dav_new | 0.23955281 | 0.08291792 | 0 |
| dav_rup | 0.56416142 | 0.20889173 | 2.33E-318 | dav_rup | 0.24297981 | 0.08341706 | 0 |
| fru_new | 0.50394301 | 0.23324547 | 0 | fru_new | 0.22201218 | 0.08579032 | 0 |
| fru_rup | 0.52722745 | 0.21385034 | 9.72E-319 | fru_rup | 0.22923039 | 0.08534064 | 0 |
| new_rup | 0.58867356 | 0.24996978 | 0 | new_rup | 0.25806691 | 0.099132 | 0 |
| ***π*** | | | | | | | |
| species | coeff | r2 | pval | species | coeff | r2 | pval |
| car | -0.0026945 | 0.00338977 | 2.91E-06 | car | 0.00102094 | 0.0004971 | 6.35E-05 |
| dav | 0.0034359 | 0.00441244 | 9.44E-08 | dav | 0.00522008 | 0.01081117 | 4.75E-78 |
| fru | 0.00287774 | 0.00296914 | 1.21E-05 | fru | 0.00458967 | 0.0075898 | 3.38E-55 |
| new | 0.00354034 | 0.00587786 | 7.17E-10 | new | 0.0039479 | 0.00740531 | 5.55E-54 |
| rup | 0.00148633 | 0.00090986 | 0.01545792 | rup | 0.00368041 | 0.00584709 | 6.07E-43 |
